# Mapping Visual Contrast Sensitivity and Vision Loss Across the Visual Field with Model-Based fMRI

**DOI:** 10.1101/2024.10.29.619403

**Authors:** Hugo T. Chow-Wing-Bom, Matteo Lisi, Noah C. Benson, Freya Lygo-Frett, Patrick Yu-Wai-Man, Frederic Dick, Roni O. Maimon-Mor, Tessa M. Dekker

## Abstract

Peripheral vision is crucial for daily activities and quality of life, yet traditional measures of visual function like visual acuity primarily assess central vision. Visual field tests can evaluate peripheral vision but require extended focus combined with precise fixation, often very challenging for patients with severe sight loss. Functional MRI (fMRI) with population receptive field (pRF) mapping offers a non-invasive way to map scotomas but is limited by its reliance on single contrast levels and the necessity of accurate fixation.

We developed an fMRI-based approach to measure contrast sensitivity across the visual field without the need for precise fixation. By combining large-field stimulation with varying spatial frequencies and contrast levels with either pRF mapping or a retinotopic atlas based on anatomical landmarks, we modeled contrast sensitivity in the primary visual cortex (V1) over a large (40 deg) expanse of the visual field. In seven normal-sighted participants, we characterized differences in V1 cortical sensitivity across eccentricities and visual quadrants, finding reliable and reproducible patterns of sensitivity differences at individual and session levels. To assess the method’s tolerance to fixation variability, we further investigated how different levels of eye movement affect cortical sensitivity patterns in two participants. We found that cortical sensitivity patterns were largely preserved across eye movement, particularly at low spatial frequencies. This suggests that our approach can accommodate several degrees of fixation instability, making it suitable for populations with unstable or biased fixation for whom visual field maps are harder to acquire behaviorally (e.g., patients with dense central scotoma or strabismus). Additionally, our method effectively visualized cases of simulated and disease-linked sensitivity loss at the cortical level. Crucially, we demonstrated that these results could be largely recovered using a structure-based retinotopic atlas, eliminating the need for pRF mapping and precise fixation - although such an approach reduced sensitivity.

This approach, integrating large-field stimulation with a retinotopic atlas, offers a promising tool for monitoring vision loss and recovery in patients with various visual impairments, addressing a significant challenge in current clinical assessments.

## Introduction

At least 2.2 billion people world-wide experience blinding eye diseases that emerge or progress across the lifespan, with 1.1 billion living with vision loss in 2020 (Bourne et al., 2021; World Health Organization, 2019). Accurate characterization of visual function is critical for optimal intervention and patient support, but fraught with challenges. The gold clinical standard measure is visual acuity, the highest spatial resolution that can be discerned. This index mainly relies on a well-functioning fovea, the small retinal area with the highest density of photoreceptors, which processes the central 0-2 degrees around fixation. However, many forms of visual impairment do not affect the fovea alone. The vast majority of our visual input comes from peripheral vision, which provides coarse but invaluable information (Alvarez, 2011; Oliva, 2005) – nearly all daily functions critically depend on intact functionality of the wider visual field, including driving, crossing the street, social function, mobility, and even reading (Lange et al., 2021). Accordingly, impairment of tissue beyond the fovea and its projections drastically affects function and quality of life (Lange et al., 2021; Lisboa et al., 2013; Roh et al., 2018; Subhi et al., 2017). Still, testing visual function across the visual field remain limited in clinical and therapeutic contexts, especially in patients with drastic central vision loss. In this study, we aimed to address this gap by introducing a novel fMRI-based approach to measure visual field sensitivity across a wide expanse of the visual field (40° diameter).

Beyond visual acuity, functional impairment across the wider visual field can be measured using a range of visual field tests, from the finger counting visual confrontation field test to more complicated and/or computerized tests (e.g., standard automatic perimetry, kinetic perimetry, microperimetry; Rai et al., 2024). Computerized tests typically involve measuring sensitivity to the luminance contrast of a target relative to a background at different visual field locations while the participant’s gaze is fixed on a central point. In some cases (e.g., microperimetry), sensitivity measurements are paired with fundus imaging, offering greater precision in linking visual field functions to specific retinal locations (Rai et al., 2024). As a result, visual field assessments can reveal functionally relevant deficits – including localized sensitivity loss and scotomas – that are not captured by foveal acuity alone, and are therefore potentially valuable for tracking disease progression and therapeutic efficacy.

Despite their clinical relevance, visual field testing comes with challenges and limitations, and as a result, the inclusion of visual field measures in sight-rescuing therapy trials is limited. Firstly, it requires prolonged fixation and sustained visual attention. This can be very challenging for patients with severe vision loss, who often struggle to fixate, and strain to detect even high-intensity stimuli. This can lead to long and unpleasant testing sessions with unreliable results. Secondly, as perception of light stimuli is inherently subjective (Rai et al., 2024) and effortful, patients may vary in their criteria for visual recognition, and in their ability to report visual signals that are weakened or distorted by disease. Together, these constraints reduce the feasibility, robustness, and interpretability of conventional visual field testing in clinical trials, underscoring the need for alternative or complementary approaches that can assess functional vision while placing fewer demands on subjective reporting.

Functional MRI (fMRI) has recently been proposed as a promising alternative to measure visual field loss, as it requires no overt task, and instead measures visual sensitivity directly from brain responses (Farahbakhsh et al., 2022; Prabhakaran et al., 2021; Ritter et al., 2019). Population receptive field (pRF) mapping fMRI can measure which parts of the cortex respond to which parts of the visual scene (Dumoulin & Wandell, 2008). This is achieved by measuring local changes in cortical blood oxygenation in response to high-contrast stimuli such as flickering checkerboards that systematically traverse the visual field. These neural timeseries are then fit with a retinotopic spatial tuning function. The fMRI signal strength in a cortical region that encodes a specific visual field location provides information about visual sensitivity in this location (Dumoulin & Wandell, 2008). This approach has recently been used to accurately recover blind spots in the visual field detected with visual field perimetry (Pawloff et al., 2023; Ritter et al., 2019). These results in turn suggest that fMRI may be a useful tool to monitor disease progression and recovery in visually impaired patients.

However, there are also significant challenges associated with this pRF mapping. Because pRF models require knowledge of where visual stimuli fall on the retina, this approach requires accurate and sustained fixation. If fixation is poor, the visual field locations assigned to cortical regions will be displaced, leading to noise or bias in the pRF maps. Additionally, most fMRI studies measure vision across a small central part of the visual fields (∼10 degrees of eccentricity) which will not capture critical dynamics in the periphery. Finally, most studies use a single maximum contrast stimulus to assess visual function (Broderick et al., 2022; Farahbakhsh et al., 2022; Liu et al., 2006; O’Connell et al., 2016; Ritter et al., 2019). This is suboptimal for at least two reasons: First, it may reduce the ability to detect subtle but clinically important changes that do not result in complete sensitivity loss or recovery. Second, with a single contrast stimulus, it becomes more difficult to determine whether a reduction in fMRI signal across sessions or brain regions reflects vision-related changes or other factors. Together, these factors currently limit the utility of neuroimaging-based visual field tests.

In this study we sought to develop fMRI methods for measuring visual field sensitivity to address these challenges, and to test its utility in artificial and realistic cases of visual field loss. Rather than presenting a single maximum contrast eccentricity-scaled checkerboard in a pRF mapping protocol, observers viewed large-field sinusoidal gratings at a range of contrasts without strict requirement to fixate. We used a large screen set up (40° diameter) to measure visual function across a large expanse of the visual field. To quantify visual sensitivity, we fit a simple cortical contrast response model to voxel timeseries evoked by the gratings, avoiding reliance on arbitrary signal amplitude thresholds. We first tested if this approach could sensitively and reliably quantify predicted contrast sensitivity differences across eccentricity and polar angle dimensions of the entire visual field (Figure 1), using a short, patient-friendly protocol in sighted controls. Critically, to achieve this we linked cortical contrast sensitivity estimates to visual field locations using retinotopic tuning estimates obtained in a separate session. However, for patients with severe vision loss it is infeasible to obtain accurate pRF measures due to poor ability to fixate a central target. We therefore also evaluated the approach using the structure-based atlas of retinotopic values developed by Benson et al. (Benson et al., 2014; Benson & Winawer, 2018). This atlas predicts retinotopic organization by aligning individual cortical anatomy (e.g., surface curvature) to a group-average template that incorporates an algebraic model of retinotopy (Benson et al., 2014). Once the subject’s brain is aligned to this structural atlas, retinotopic maps defined by the model – i.e., polar angle and eccentricity maps – are projected onto the individual’s cortex. This allows estimation of visual field maps without requiring functional imaging, and provides a non-invasive, anatomy-driven approximation of visual field representations. To increase accuracy of the Benson atlas for our large-field stimuli, we subsequently fit these eccentricity estimates to the Horton and Hoyt (1991) model of cortical magnification in each individual.

**Figure 1.**
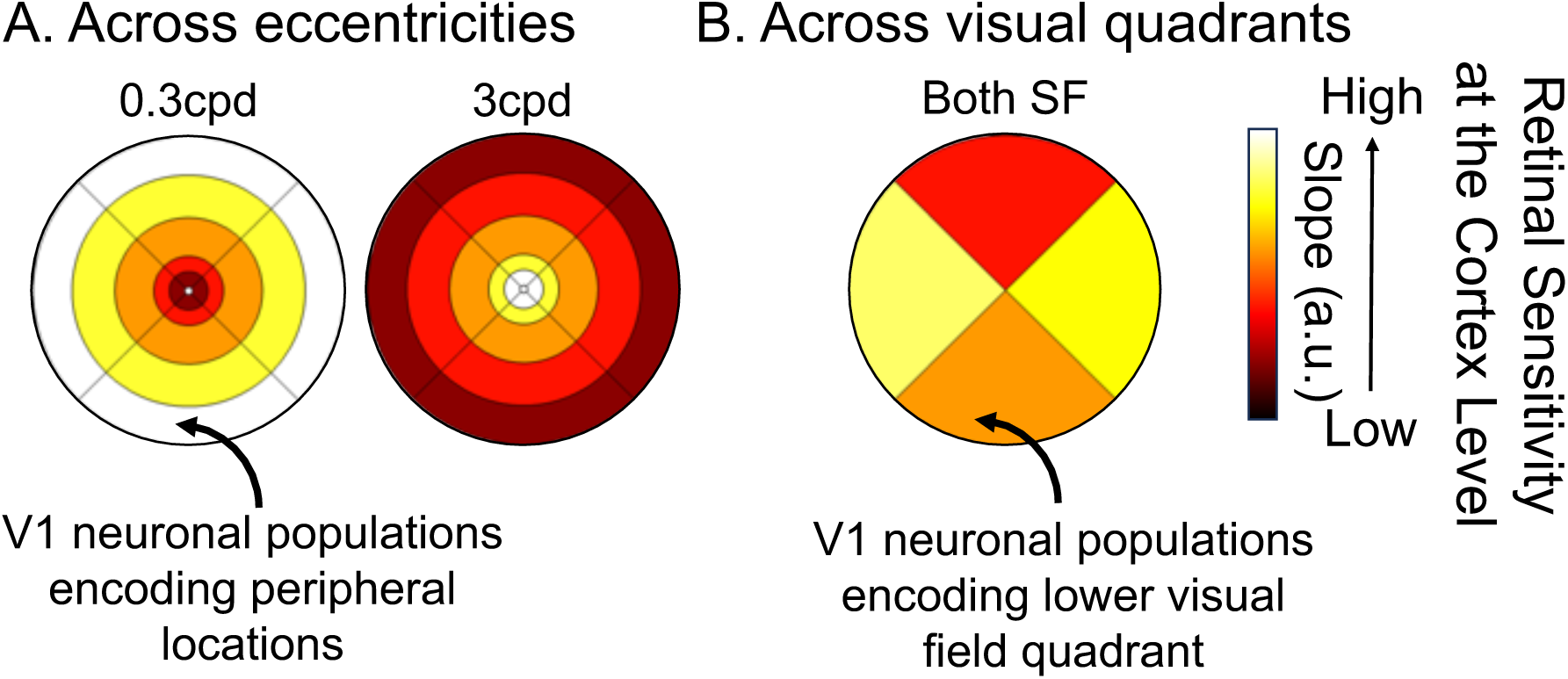
Expected changes in cortical contrast sensitivity in typically sighted controls for two different spatial frequencies (low spatial frequency, 0.3 cycles per degree (cpd), and high spatial frequency, 3.0 cpd), A: as a function of eccentricity and B: visual field quadrants. With reference to eccentricity, we expect higher sensitivity of V1 neuronal populations to 1) lower spatial frequencies in the periphery, and 2) higher spatial frequencies in the center of the visual field. By comparison, we expect both high and low spatial frequency stimulation to evoke greatest sensitivity in V1 neuronal populations encoding the left and right visual quadrants, less in the lower, and least in the upper quadrant.

We demonstrate that our approach can recover, at a cortical level, known sensitivity changes across the visual field. V1 neuronal populations encoding the central visual field show the highest sensitivity to high spatial frequencies, whilst those encoding the periphery are most sensitive to low spatial frequencies. We also find greater sensitivity in V1 regions receiving visual inputs from the horizontal compared to vertical quadrants, and from the lower compared to upper quadrants. Crucially, these effects are detectable at the individual level and repeatable across visits, suggesting our approach will be suitable for detecting visual field sensitivity changes over time (e.g., due to learning, ageing, disease progression, or recovery). Moreover, when using the Benson atlas instead of individual pRF measures to link neuronal responses to visual field locations, we still detect these effects, although with slightly reduced sensitivity. We also demonstrate the clinical relevance of this approach by recovering simulated scotomas (i.e., a ring of visual field loss around fixation and the loss of an entire visual field quadrant), as well as visual field loss in a patient with a neurodegenerative disorder causing large areas of visual field loss. We conclude that this approach, i.e., the combination of cortical contrast sensitivity testing, large field stimulation, and a large-field calibrated retinotopic atlas, may be particularly useful for quantifying visual recovery in patients, whether spontaneous or following gene-therapy interventions, and for linking brain-based activations to behavioral performance.

## Methods

### Participants

We collected data from 7 healthy controls (mean±SD: 29.6±4.7yo; 1M). All controls either had normal or corrected to normal vision, with no other ocular pathologies, and were recruited from the local staff and student pool at the University College of London. Each control completed both the population receptive field (pRF) mapping and the fMRI contrast sensitivity task. To assess measurement repeatability, four participants (C2, C4, C5, C6) performed the contrast sensitivity task twice. Additionally, one participant (C5) repeated the task under two simulated vision loss conditions (ring or quadrant loss), and two others (C5, C6) completed it with different levels of eye movement.

We also recruited a patient with Leber Hereditary Optic Neuropathy (M, 45.5 years old, mutation: m.11778G>A, disease duration: 36.1 years), who presented with relatively stable fixation and was able to complete both the pRF mapping and the fMRI contrast sensitivity task. All participants signed a written informed consent to participate in the study, and this study was approved by HRA/HCRW and Research Ethics Committee (IRAS 258959; REC 20/YH/0336).

### Stimulus & Apparatus

Stimuli were programmed in MATLAB 2020b (MathWorks, Natick, Massachusetts, USA) using the Psychtoolbox-3, and displayed using an EPSON LB-1100U projector (resolution: 1920x1200 pixels, 60Hz, projected area: 43.2x27cm). Because the projector was too bright and to reduce discomfort in the scanner, a circular neutral density filter (ND4 Plus+; Urth, Byron Bay, Australia) was placed in front of the projector to reduce light intensity by 4 times without altering the colors. The projector was also gamma-corrected to ensure accurate representation of the intended contrast levels and spatial frequencies used in our experiments. In all tasks, stimuli were embedded in circular aperture of 42.3x42.3° diameter. To achieve the large field-of-view, the front part of the 64-channel MRI coil was removed, and stimuli were back-projected onto a screen positioned close to the participant’s head inside the bore (see Figure A1 in *Appendix* section). Participants viewed the stimuli through a mirror, for a total viewing distance of 34cm. Throughout the session, participants were constantly reminded to remain still to minimize head and body motion, and eye motion and alertness were monitored using an EyeLink 1000Plus device (SR Research, Ottawa, Canada). In all tasks, participants were asked to report when the color of a central fixation dot changed.

### fMRI Contrast Sensitivity Task

Participants were presented binocularly with achromatic sine-wave gratings, flickering at a temporal frequency of 2Hz and varying in contrast levels and spatial frequencies. Contrast levels were defined as Michelson contrast 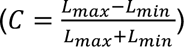, where Lmax and Lmin are maximum and minimum luminance. Each stimulus, presented for 13s, featured a specific contrast level (either 7.5, 42.2, 60, or 100%) and spatial frequency (either 0.3 or 3 cycles per degree). Stimuli were then followed by 2s of a grey background (0% contrast) to minimize the experience of after-effects. Each combination of contrast level and spatial frequency was shown three times within a run, arranged in a pseudo-random sequence, for a total of 9 repetitions per combination (i.e., 3 runs). Eight additional blocks of 15-second rest periods with a grey background were introduced to allow sufficient baseline. Four participants (C2, C4, C5, C6) were invited for a second session in which they repeated the task to assess the reliability of the measures.

For six control participants (C1–C6), gratings were initially presented with a fixed horizontal orientation. In an updated version of the task – used for C7, cases of simulated eye movement, cases of artificial scotoma, and the LHON patient – the orientation varied every 5 s among four angles (−45°, 0°, 45°, 90°). Contrast sensitivity patterns were consistent across single- and multiple-orientation conditions, including in participants who completed both versions, indicating robustness across orientation-tuned populations.

#### Effect of eye movement

Participants C5 and C6 also performed a version of the task designed to test the effect of eye movements. In this version, saccades were elicited by randomly and rapidly shifting the fixation dot away from central fixation (C5: 2° and 5° from fixation and random motion; C6: up to 2° from fixation). Participant C5 was tested using 0.3 and 3cpd gratings at four contrast levels (7.5, 42.2, 60, 100%), while participant C6 was tested only under the low spatial frequency condition (0.3cpd).

Fixation stability was assessed for each fMRI run using the bivariate contour ellipse area (BCEA), which estimates the area (in degrees² or arcmin²) of an ellipse that contains approximately 95% of fixation points. BCEA was calculated using the formula: *BECA* = 2 ∗ *k* ∗ 𝜋 ∗ 𝜎*_h_* ∗ 𝜎*_v_* ∗ 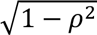, as described by Morales et al. (2016). In this expression, 𝜎*_h_* and 𝜎*_v_* represent the standard deviations of eye position in the horizontal and vertical directions, respectively, while 𝜌 corresponds to the Pearson correlation coefficient between horizontal and vertical eye positions. The constant *k* determines the size of the ellipse based on the desired probability area, defined by the relationship 𝑃 = 1 – 𝑒^−*k*^, with P set to 0.95 in this study. A smaller BCEA indicates greater fixation stability.

#### Simulated vision loss

One healthy control participant (C5) also performed a version of the task designed to simulate two forms of visual input loss (i.e., artificial scotoma). These simulations were implemented by: (a) masking a region of the visual field with a grey, annular ring, covering 3°-8° eccentricity, and (b) masking the upper-right visual quadrant using a grey quarter-sector overlay. The stimuli and contrast levels used in this task were identical to those described in the original task.

### Population Receptive Field (pRF) mapping

To investigate how distinct regions of the visual field map onto the cortex, we collected pRF mapping data using a standard ring-and-wedge stimulus (Dumoulin & Wandell, 2008). Participants were presented binocularly with a black-and-white checkerboard, phase-flickering at a temporal frequency of 2Hz, and at maximum contrast level. The stimulus spanned up to the boundaries of the circular aperture (i.e., 42.3x42.3° diameter) and was scaled across eccentricity to account for cortical magnification. This scaling was implemented by applying a logarithmic transformation of retinal radius, such that radial checker boundaries were defined in log-eccentricity space (𝑙𝑜𝑔(𝑟), where 𝑟 denotes to eccentricity relative to the fixation target). This produced an exponential increase in checker size with eccentricity (scaling factor = 3.2; ∼1.37 times increase per radial step), resulting in lower spatial-frequency content at larger eccentricities – consistent with known variations in V1 spatial-frequency tuning. Because this eccentricity-dependent scaling assumes precise fixation, it can be challenging for individuals with central vision loss, further motivating the use of Benson atlas templates in such populations. Three runs of 352 volumes were collected per participant.

### MRI acquisition

Data were collected at the Birkbeck-UCL Centre for NeuroImaging (BUCNI, London, UK), on a Siemens PRISMA 3 Tesla scanner.

Functional data were acquired using a 64-channel head coil with the front part removed, to allow an unobstructed view of the screen whilst ensuing enough signal from the remaining 40 effective channels covering the side and back of the head. T2*-weighted echo-planar imaging were collected with an accelerated multiband sequence kindly provided by CMRR (version R016a, https://www.cmrr.umn.edu/multiband; Cauley et al., 2014; Xu et al., 2013); multi-band factor: 4, voxel resolution: 2mm isotropic, FOV: 212x212x96mm, flip angle: 60°, repetition time (TR): 1000ms, echo time (TE): 35.2ms, echo spacing: 0.56ms, bandwidth: 2620Hz/Px, with 48 transverse slices angled to be approximately parallel to the calcarine sulcus whilst avoiding the orbital cavities). The same sequence was used to acquire 4 additional scans in the opposite phase-encoding direction, to allow B0 deformation correction in the pre-processing steps.

A T1-weigthed structural image was acquired with a 32-channel coil, using MPRAGE sequence (voxel resolution: 1mm isotropic, 208 slices, FOV: 256 x 256 x 208mm, flip angle: 7°, TR = 2300ms, TE = 2.98ms, TI = 900ms, bandwidth: 240Hz/Px, echo spacing: 7.1ms, acquisition time: 5min30s).

### Data analysis

Segmentation of the anatomical scan and reconstruction of cortical surfaces were performed in FreeSurfer 7.1.1.

Functional data were pre-processed using a combination of AFNI 24.1.22, FreeSurfer and FSL 6.0.7.12 commands. The following steps were followed for both the contrast sensitivity task and the pRF mapping task. First, initial correction for distortions caused by field inhomogeneity was performed using a blip-up/blip-down approach (AFNI’s unWarpEPI.py program), and the first 4 volumes of each run were discarded using FSL’s fslroi program. Then an alignment scan was created from the volume out of all collected runs that contained the least amount of voxel outliers using AFNI’s 3dToutcount, 3dTstat and 3dcalc programs. This alignment scan was then co-registered to the MPRAGE image using FreeSurfer’s bbregister program, resulting in a rigid-body transformation matrix. In case of misregistration, the alignment scan was defined as one of the single-band reference scans, before re-running the co-registration step. Motion correction was then carried out by aligning each functional volume from all runs to the previously defined alignment scan using AFNI’s 3dvolreg program.

### Processing of the contrast sensitivity task

To accurately capture neural activity across various eccentricities and polar angle locations, minimal smoothing (0.5mm FWHM Gaussian blur) was applied to the contrast sensitivity task data using FSL’s 3dmerge program. This was done to meet the minimum requirements of the GLM module in SPM. The contribution of each contrast/spatial frequency combination to the BOLD signal was modelled using the general linear model (GLM) approach in SPM12 (https://www.fil.ion.ucl.ac.uk/spm/), with motion parameters as additional regressors. This step resulted in volumetric images of *β*-values, each reflecting the specific contribution of a given contrast/spatial frequency combination to the baseline BOLD signal. These resulting 𝛽-images were then projected onto the individual’s cortical surface using FreeSurfer’s mri_vol2surf program, with no surface smoothing and the default fractional projection along surface of 0.5. The latter sets the fraction (0,1) of the cortical thickness at each vertex to project along the surface. Surface-projected 𝛽-images were finally converted into a MATLAB-compatible format for further analysis using SamSrf’s samsrf_mgh2srf program.

### Processing of pRF mapping task

Pre-processed volumes from the pRF mapping task were projected onto the cortical surface using FreeSurfer’s mri_vol2surf program. Here, we used the previously generated rigid-body transformation matrix to ensure accurate spatial alignment and smoothed on the cortical surface with a Gaussian kernel with a FWHM of 3mm.

During pRF model fitting, pRF centers were allowed to extend beyond the stimulated visual field to improve model stability near stimulus boundaries – up to approximately 1.5 times the maximum stimulus eccentricity (∼30°). Eccentricity was sampled on a logarithmically spaced grid defined as 2^)^, with 𝑥 ranging from −5 to 0.6 in steps of 0.2, and then scaled by the maximum stimulus eccentricity (20°) to express pRF centers in degrees of visual angle. This sampling scheme provided finer resolution near the fovea and progressively coarser sampling at larger eccentricities, consistent with cortical magnification principles.

The resulting outputs were then converted into a MATLAB-compatible format for further analysis. pRF estimates were computed by fitting a 2D Gaussian model to the data, using the SamSrf toolbox (version 7.13; https://github.com/samsrf), resulting in three pRF parameters for each vertex on the cortical surface: the x- and y-coordinates of the pRF center, and the pRF size (σ). Eccentricity and polar angle maps were computed from the x- and y-coordinates.

### Regions-Of-Interest

In this study, we were interested in how visual information across the visual field is processed in early visual areas, notably in the primary visual cortex V1. Early visual regions were manually delineated for each participant in SamSrf v7.13, based on the output of the pRF fitting procedure and standard functional criteria to identify borders between V1, V2, and V3 (DeYoe et al., 1996; Engel, 1997; Sereno et al., 1995). In this report, we focused our interest on V1, given its crucial role in the initial processing of basic visual features like spatial frequency and contrast. For all subsequent analyses of cortical contrast sensitivity, pRF centers outside the stimulated 20° eccentricity were excluded. Similarly, although the Benson atlas provides eccentricity estimates extending far beyond the stimulated range (up to ∼90°), only values within 20° were retained to maintain consistency across pRF-based and atlas-based analyses.

### Quantifying eccentricity- and location-dependent variations in sensitivity

#### Using subject-specific pRF estimates

At this stage, for each vertex on the cortical surface, we obtained 𝛽-values for each tested contrast/spatial frequency combination in the contrast sensitivity task, as well as pRF parameters resulting from the fitting of the pRF model to the pRF data (Figure 2). 𝛽-values from the contrast sensitivity task in each vertex were then filtered based on ROI location (V1) and pRF parameters (R^2^≥0.05; pRF size: 𝜎≤6).

**Figure 2.**
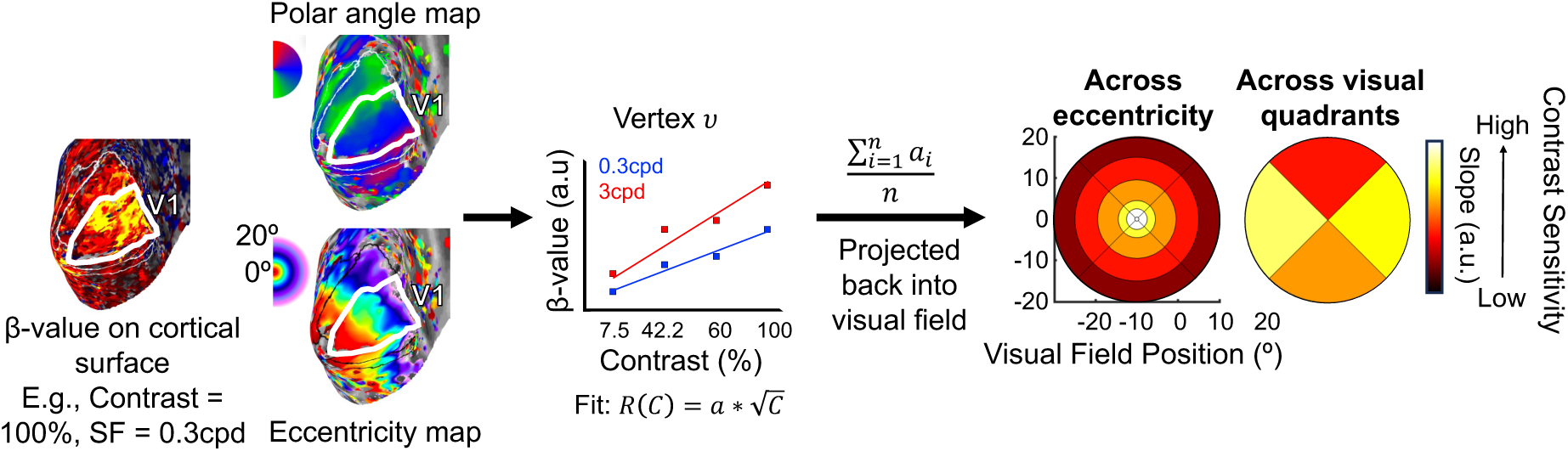
Estimation of cortical contrast sensitivity across the visual field. Participants performed two fMRI tasks in the scanner: a contrast sensitivity task and a pRF mapping task. For each individual and tested spatial frequency, the contribution of each stimulus contrast level to the BOLD signal in the contrast sensitivity task was estimated using a GLM approach. Resulting β-coefficients for each contrast level were then projected onto the individual’s cortical surface. For each vertex in V1, a square root function (𝑅(𝐶) = 𝑎 ∗ √𝐶) was then fitted to these data, taking the estimated slope *a* as measure of cortical contrast sensitivity. V1 slopes were finally averaged across vertices based on their eccentricity and polar angle preference as estimated with pRF mapping, and across participants. Mean slopes were then projected back into the visual space, producing a heatmap projection of V1 contrast sensitivity across eccentricities and along visual field quadrants.

To assess contrast sensitivity, we modelled the neural response (i.e., 𝛽-values) in V1 as a linear function of the square root of the contrast values (𝑅(𝐶) = 𝑎 ∗ √𝐶). This model rests on the assumption of a monotonic relationship between the presented stimulus contrast level and the resulting BOLD response and has been used in previous work by Buracas and colleagues to investigate contrast sensitivity at the cortex level in healthy controls (Buracas et al., 2005; Buracas & Boynton, 2007). Steeper slopes within this context indicate greater V1 contrast sensitivity.

We fit this model for each vertex separately, for each participant and spatial frequency (0.3 and 3cpd) condition. The intercept was constrained to be zero, as the distribution of *β*-values is expected to be centered on zero when the contrast is zero (i.e., when no stimulus is presented). Next, the resulting V1 slope estimates were binned and averaged over five regions using pRF eccentricity estimates: 0.5°-2.5°, 2.5°-4.5°, 4.5°-9.5°, 9.5°-15°, and 15°-20°. These bins relate to the clinical definition of visual space based on anatomical landmarks, corresponding to the fovea, parafovea, perifovea and two near-periphery regions, respectively. Similarly, slope estimates were spatially constrained within a ±45°-wedge centered on each cardinal meridian to investigate asymmetries as a function of pRF polar angle locations, thus dividing the visual space between the upper, lower, left and right quadrants. These wedges were also derived from the polar angle estimates obtained with pRF mapping.

For each participant, average slopes for each eccentricity bin and polar angle wedge were then projected back into their corresponding visual field locations. This resulted in heat-maps of V1 contrast sensitivity across eccentricities and visual field quadrants, for each tested spatial frequency condition at an individual level (Figure 2). Finally, average slopes for each eccentricity and polar angle bin from each participant were used to compute the group-level average.

#### Using calibrated Benson retinotopic template atlas

The same approach was used to quantify changes in V1 contrast sensitivity across the visual field, using a retinotopic template (Benson et al., 2014; Benson & Winawer, 2018) instead of pRF mapping. This template was fitted to each subject’s anatomical MPRAGE image using the Python library neuropythy (Benson & Winawer, 2018), which provided automatic delineation of early visual regions, based on FreeSurfer’s anatomical alignment. Additionally, it offered eccentricity and polar angle estimates for each vertex on the cortical surface. The advantage of this method over pRF mapping is that it does not require fixation or pRF scanning, making it a valuable tool to use when scan time is valuable or in populations where good fixation is hard to achieve.

We initially observed inaccuracies between the template and individual retinotopy eccentricity estimates which led to substantial distortions in cortical visual field maps due to cortical magnification – especially in peripheral locations (see Figure A4 in *Appendix* section). To enhance the accuracy of eccentricity estimates in the template map, we aligned the distribution of eccentricity values from the fitted Benson template atlas with Horton and Hoyt’s model (Horton & Hoyt, 1991; Figure 3A). This model suggests that the linear cortical magnification factor (𝑀*_linear_* in mm/°) is inversely proportional to eccentricity (𝐸 in degrees), expressed as 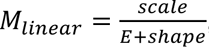, with the normalizing parameter 𝑠ℎ𝑎𝑝𝑒 equal to 0.75°, as measured by Horton and Hoyt (1991). Using this relationship, we first estimated for each hemisphere the *scale* value based on each individual’s V1 surface area (from the Benson V1 label) and the minimum and maximum eccentricities considered in the model (0° and 90°). The mean *scale* value for the left and right V1 hemispheres across all tested controls were 15.0±0.99 and 16.0±1.09, respectively. In comparison, Horton and Hoyt (1991) measured a mean *scale* of 17.3 mm. The mean V1 surface areas for the left and right hemispheres were 2700.7±358.7mm^2^ and 3067.7±416.3mm^2^.

**Figure 3.**
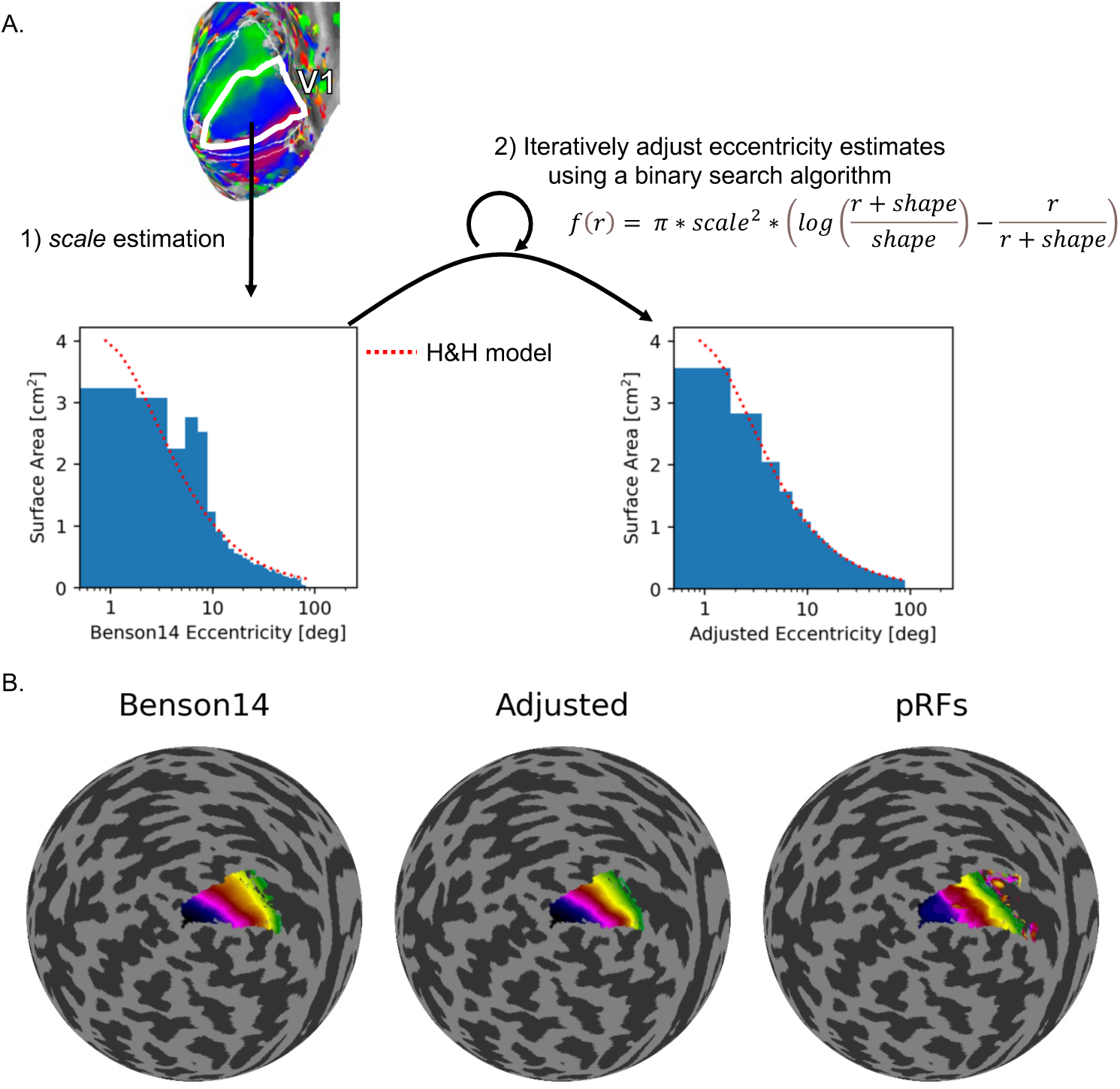
Eccentricity alignment and surface area estimation in V1. A: For each hemisphere, the scale parameter in Horton and Hoyt’s linear magnification model 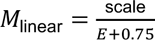 was estimated from individual V1 surface area and the 0°-90° eccentricity range. Using this, we derived a cumulative surface-area function 𝑓(𝑟), which gives the total cortical area of V1 representing eccentricities from 0° to 𝑟. For visualization, the histograms show the differential form of this function – i.e., the surface-area density as a function of eccentricity. This corresponds to the derivative 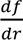, which reflects the areal magnification (denoted 𝑚∗(𝑟)) integrated over polar angle. This representation provides a clearer view of how cortical surface area is distributed across eccentricities while remaining consistent with the underlying cumulative function 𝑓(𝑟). Adjusted eccentricity values were obtained by iteratively matching the empirical surface-area distribution to the model using a binary search procedure. B: Comparison of eccentricity maps generated using the Benson template, the adjusted eccentricity values, and population receptive field (pRF) estimates, showing improved alignment with empirical data.

We then defined a function 𝑓(𝑟) that describes how the surface area of V1 changes with eccentricity (Equation 1; Figure 3A). This was achieved by integrating Horton and Hoyt’s function over the visual field, resulting in an equation that calculates the amount of V1 surface area from the foveal confluence up to a particular eccentricity (𝑟). Next, we sorted the input eccentricity values and calculated the cumulative surface areas. A numerical method was then employed to find the eccentricity values that match the desired distribution. This method involved iteratively adjusting the range of possible values until the calculated values closely matched the desired values. Finally, the adjusted eccentricity values were returned in their original order and values from both hemispheres were merged to provide bilateral eccentricity estimates.

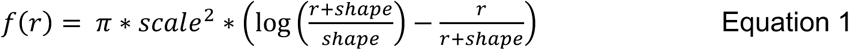

## Results

To measure visual field function, we developed a new measure of cortical contrast sensitivity, assessing the brain’s ability to discriminate gratings of varying spatial frequencies based on luminance variations. This depends on many factors, including the size, orientation, and density of cells in different parts of the retina. As these factors change across the visual field, contrast sensitivity varies with spatial frequency and visual field location in the typical, healthy visual system. We first investigated whether our brain-based approach can reliably detect these known differences in contrast sensitivity as a function of eccentricities and visual field quadrants, in individual normal sighted controls. We then investigated whether our approach could recover loss of visual inputs at the cortical level, either simulated in a healthy control or pathological in a patient. In each section, findings were first reported for the pRF-defined ROIs and then for the Benson-defined ROIs.

### Spatial frequency preference changes in V1 neuronal populations across eccentricities

For high spatial frequency stimuli, we expected better sensitivity in V1 neuronal populations encoding central locations in visual space than in those encoding the periphery. In contrast, for low spatial frequency information, we expected the opposite, better sensitivity in V1 neuronal populations encoding the periphery than the center of the visual field.

Using ANOVA, we found the expected interaction between spatial frequency and eccentricity (𝐹(1.96,11.79) = 28.66, p < 0.001; Figure 4) as well as a main effect of eccentricity (𝐹(2.33,13.99) = 12.67, p < 0.001). These effects were statistically significant at the individual participant level showing high sensitivity (Figure 4A, transparent lines show individual data; Figure 4C shows individual visual field plots; for stats, see Table A1 in *Appendix* section).

**Figure 4.**
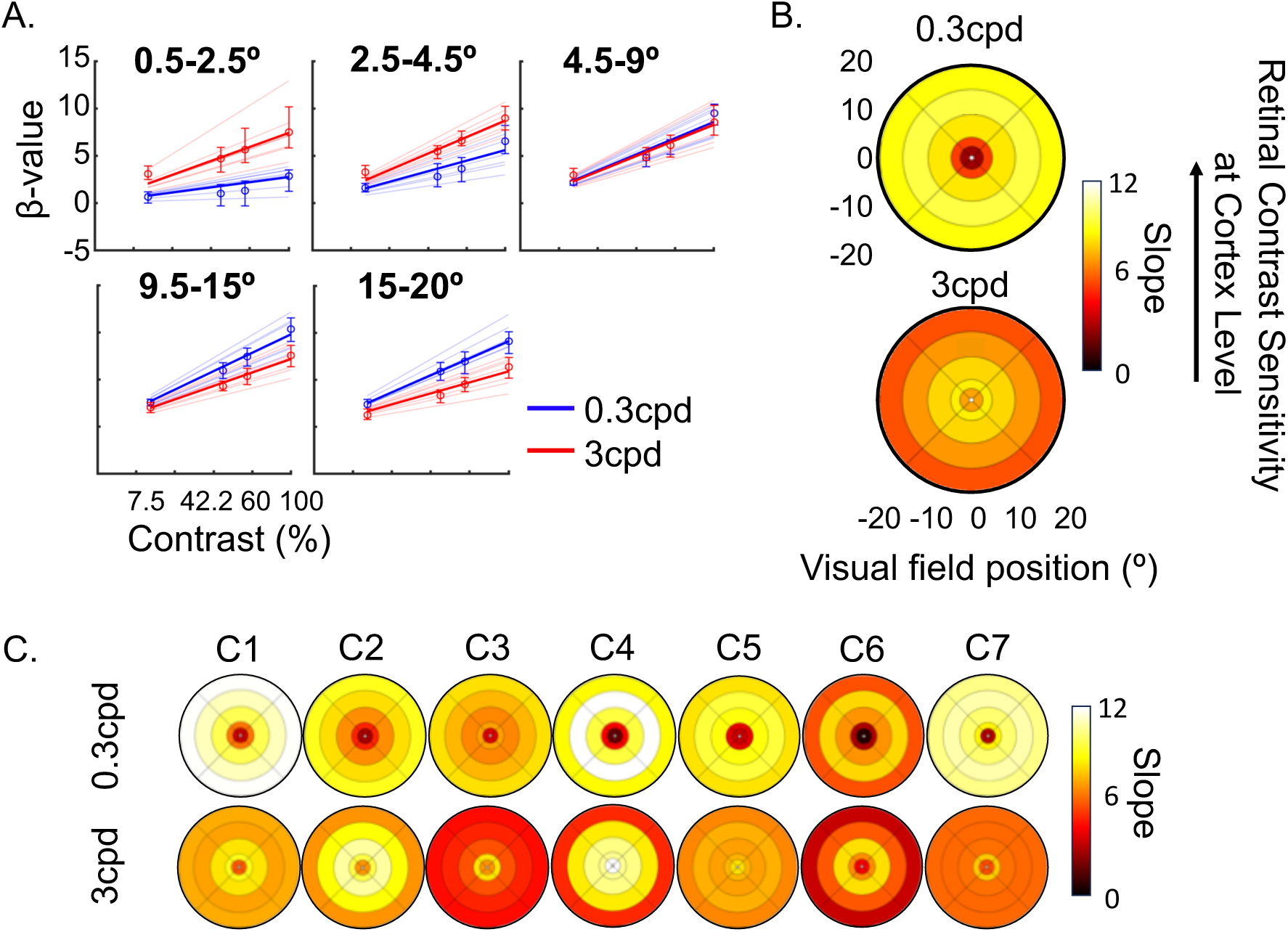
Spatial frequency preference V1 across the visual field. For each participant, five eccentricity bins were defined using subject-specific pRF estimates: 0.5°-2.5°, 2.5°-4.5°, 4.5°-9.5°, 9.5°-15°, and 15°-20°. **A:** Average β-values versus stimulus contrasts for low (0.3cpd; blue) and high (3cpd; red) spatial frequencies across eccentricity bins. Error bars are the 95% confidence intervals. Thick lines represent the group-level model fit, thin lines are the individual fits. **B:** Slopes projected into visual space for low (0.3 cpd) and high (3 cpd) spatial frequencies. The color scale corresponds to slope estimates, with higher values indicating higher cortical contrast sensitivities. **C:** Heat plots for each participant (C1-C7).

Post-hoc tests using Holm-Bonferroni correction show that V1 neuronal populations receiving inputs from the central visual field (0.5-4.5°) showed greater contrast sensitivity to high spatial frequency as compared to low spatial frequency stimuli (steeper slope for the 3cpd versus 0.3cpd condition: **0.5-2.5°**: 𝑡(6) = 4.35, 𝑝_45,6_ = 0.0149; **2.5-4.5°**: 𝑡(6) = 3.471, 𝑝_45,6_ = 0.0266). Conversely, peripheral eccentricities in V1 (above 9.5°) showed higher contrast sensitivity to low as compared to high spatial frequency stimuli (steeper slope for 0.3cpd versus 3cpd condition: **9.5-15°**: 𝑡(6) = −4.591, 𝑝_45,6_ = 0.0149; **15-20°**: 𝑡(6) = −6.615, 𝑝_45,6_ = 0.0029). Between 4.5° and 9.5°, V1 contrast sensitivity was similar for both spatial frequencies (𝑡(6) = −0.226, 𝑝_45,6_ = 0.8286). Crucially, these effects remained when using retinotopic estimates based on structural scans derived from the Benson retinotopic atlas instead of the pRF-mapping measures (**0.5-2.5°**: 𝑡(6) = 5.768, 𝑝_45,6_ = 0.0059 ; **2.5-4.5°**: 𝑡(6) = 2.531, 𝑝_45,6_ = 0.0892 ; **4.5-9.5°**: 𝑡(6) = −0.293, 𝑝_45,6_ = 0.7792; **9.5-15°**: 𝑡(6) = −3.274, 𝑝_45,6_ = 0.0509; **15-20°**: 𝑡(6) = −3.528, 𝑝_45,6_ = 0.0496; see Figure A2 and Table A3 in *Appendix* section).

These results demonstrate that our approach can detect subtle changes in visual sensitivity across eccentricities at the individual participant level. The ability to reveal these gradients was made possible by the large peripheral coverage provided by our large-field stimulation set-up (see Figure A1 in *Appendix* section), which enabled a more complete characterization of V1 sensitivity across the visual field. Importantly, the same effects were preserved when using retinotopic estimates derived from structure-based atlases, demonstrating that atlas-based methods can be used as alternative to pRF mapping in cases where it might otherwise be difficult or impossible to directly collect pRF measures. Together, these highlight both the validity of our approach and its potential to broaden the scope of visual neuroscience.

### Contrast sensitivity is not equal across the visual field quadrants in V1 neurons

Contrast sensitivity is known to vary across the quadrants of the visual field; Previous behavioral studies have shown greater contrast sensitivity along the horizontal versus vertical meridians, and along the lower versus upper vertical meridian (Barbot et al., 2021; Carrasco et al., 2022; Himmelberg et al., 2020, 2023). Recent fMRI studies have also shown that these asymmetries in BOLD response amplitude to single contrast gratings (Kurzawski et al., 2022; Liu et al., 2006; O’Connell et al., 2016). Therefore, we expect our fMRI approach to uncover these established asymmetry patterns at the cortical level.

Figure 5 shows how cortical contrast sensitivity in V1 varies across visual field quadrants, for the 0.3cpd and 3cpd conditions. For this analysis, we collapsed eccentricities (0.5° to 20°) into a single bin. We found a main effect of visual field quadrant location on V1 sensitivity (𝐹(2.46,14.76) = 20.71, 𝑝 < 0.001). Post-hoc pairwise comparisons using Holm-Bonferroni corrections revealed that, as predicted, the cortical contrast response function had a higher slope – indicating better V1 sensitivity – along the horizontal versus vertical quadrants (Horizontal-Vertical Anisotropy – **HVA:** 𝑡(6) = 5.908, 𝑝_45,6_ = 0.0031) and along the lower versus upper quadrant (Vertical Meridian Anisotropy – **VMA:** 𝑡(6) = 4.106, 𝑝_45,6_ = 0.0126). Conversely, no difference in cortical contrast sensitivity was found between V1 neuronal populations encoding the left and right quadrants of the visual field (Left-Right Horizontal Meridian Anisotropy – **LRHMA:** 𝑡(6) = 0.7197, 𝑝_45,6_ = 0.4988). Moreover, there was no interaction between spatial frequency and visual field quadrant positions (𝐹(2.16,12.99) = 1.34, 𝑝 = 0.298), suggesting V1 visual field anisotropies are relatively constant across spatial frequencies. Importantly, all these differences are also present in individual participants (Figure 5A, transparent lines are individual data, Figure 5C shows individual visual field plots; for stats, see Table A2 in *Appendix* section). All participants had better V1 sensitivity in the horizontal versus vertical quadrants. 6 out of 7 participants showed asymmetries between the lower and upper quadrants. Differences in V1 sensitivity between left and right quadrants were inconsistent across participants (3 participants: left>right; 2 participants: right>left; 2 participants: non-significant), in line with no difference at the group-level. This again demonstrates the high sensitivity of our approach for identifying and quantifying sensitivity changes across the visual field, at the individual participant level, offering a potential valuable tool for assessing and addressing visual field loss in clinical populations.

**Figure 5.**
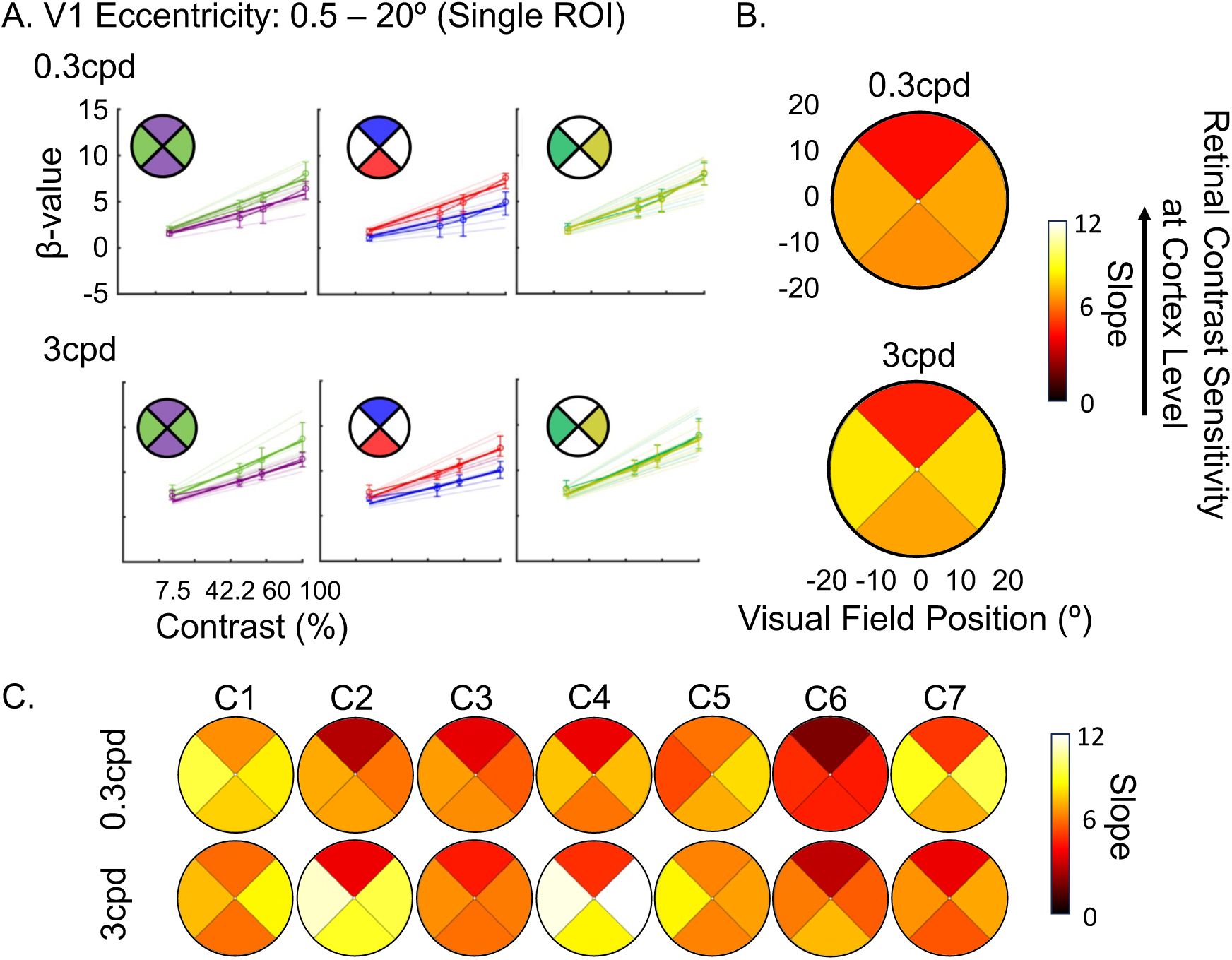
Anisotropies in V1 across visual field quadrants for low (0.3cpd; top row) and high (3cpd; bottom row) spatial frequencies. pRF estimates were used to link brain responses to visual field positions. Visual quadrants are ±45° around the cardinal meridians, with slope values for eccentricities between 0.5°-20°. Horizontal quadrants are left and right; vertical quadrants are upper and lower. **A**: Average β-values versus contrast levels for each anisotropy (horizontal vs. vertical, upper vs. lower, left vs. right). Error bars are 95% confidence intervals. Thick lines correspond to average model fit, which thin lines are individual fits. **B**: Slope values projected onto visual field quadrants. The color scale represents slope estimates, with higher values indicating greater V1 contrast sensitivity. **C:** Heat plots for each participant (C1-C7).

We next tested if we could recover these V1 anisotropies when using retinotopic estimates based on structural scans fit with the Benson atlas instead of pRF measures (see Figure A3 and Table A3-4 in *Appendix* section). We found that the horizontal-vertical anisotropy effect was recovered (**HVA:** 𝑡(6) = 3.584, 𝑝_45,6_ = 0.0347), but that the vertical meridian anisotropy effect was not (**VMA:** 𝑡(6) = 0.744, 𝑝_45,6_ = 0.9697) with this approach. These results provide evidence that our approach can still detect subtle visual field quadrant anisotropies using a structure-based retinotopic atlas, although less accurately.

While the most pronounced contrast sensitivity difference along visual field quadrants (horizontal versus vertical quadrants) was retained when the adjusted Benson atlas was used to replace pRF measurements, not all anisotropies survived. This may be because the template does not model cortical magnification differences along meridians, reducing accuracy. It is not surprising that retinotopic estimates based on structure-retinotopy relationship modeling, provides less accurate estimates of retinotopic tuning of neuron population in V1 than direct measures of the individual retinotopy. Any individual deviations from the model make it more challenging to accurately assign cortical contrast sensitivity to visual field locations, potentially decreasing the ability to detect changes in visual function across the field. Given this challenge, results with the adjusted Benson map are surprisingly similar to those based on pRF measurement, showing promise for clinical applications when pRF mapping is unfeasible.

### V1 sensitivity differences across the visual field are reproductible across visits

Having shown that changes in cortical contrast sensitivity across the visual space can be detected with fMRI, it is crucial to determine how reliable our measure is. This is especially important for potential applications in monitoring changes over time. Using Spearman correlations, we measured the reliability of cortical contrast sensitivity estimates for the 0.3cpd spatial frequency condition across two repeated measurements, collected on two different days. We collected test-retest reliability measures from 4 out of 7 participants (Figures 6A-B) and benchmarked them against the correlations between the 0.3cpd condition and 3cpd spatial frequency condition, collected in the same session (Figure 6C). If measures are reliable, correlations should be higher for repeated measures with the same spatial frequency stimulus, collected on different days. We tested this prediction using a one-tailed paired t-test. Our findings clearly show that the gradual increase in cortical sensitivity toward the periphery for the 0.3cpd condition is consistent across the two sessions (Figure 6A). Moreover, the data of individual participants show characteristic profiles that stay highly consistent. Additionally, we observed moderate-to-strong positive Spearman correlations across sessions for the 0.3cpd condition (Figure 6B; C2: 𝜌 = 0.5793; C4: 𝜌 = 0.6996; C5: 𝜌 = 0.4059; C6: 𝜌 = 0.7677), which were consistently higher than those across different spatial frequency conditions (Figure 6C; C2: 𝜌 = 0.5503; C4: 𝜌 = 0.2318; C5: 𝜌 = 0.2158; C6: 𝜌 = 0.4993). This difference was statistically significant (𝑡(3) = 2.62, 𝑝 < 0.0395). This indicates that within participants, our measure is reliable across sessions and distinguishes effectively between patterns elicited by various spatial frequency conditions. Collectively, this suggests that our measure could serve as a robust tool for detecting changes in cortical sensitivity, such as those occurring over time or following therapeutic interventions.

**Figure 6.**
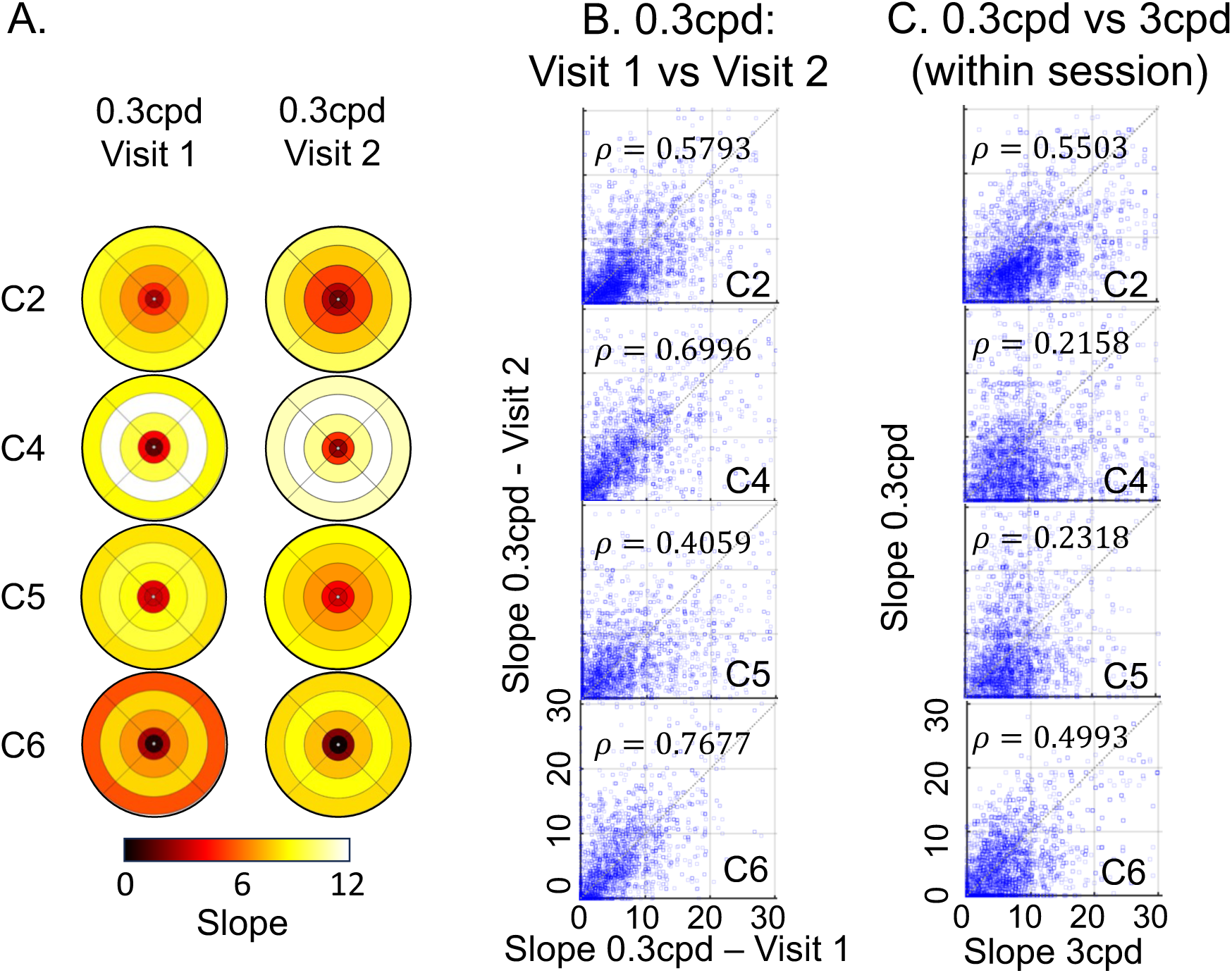
Reliability of V1 slopes across sessions, in 4 of the 7 controls. A: V1 contrast sensitivity (i.e., slopes) plotted in visual field space (using pRF estimates), showing a consistent session-independent increase in cortical sensitivity toward peripheral locations for the 0.3cpd condition. Higher slopes (warmer colors) indicate higher V1 sensitivity. B: Slopes for the 0.3cpd condition across two visits. C: Slopes for the 0.3cpd and 3cpd conditions collected within the same session.

### Effect of eye movements on V1 cortical sensitivity

So far, we have demonstrated that our measure of cortical sensitivity can reliably recover known gradients in sensitivity across eccentricities and visual quadrants. We also showed that this measure was consistent across visits and sessions, suggesting its potential utility for monitoring changes over time. However, all prior tasks were conducted under conditions of central fixation, with participants instructed to maintain gaze on a central dot. A key motivation for this approach was its theoretical robustness to fixation instability. We therefore also aimed to investigate how varying degrees of eye movement might influence cortical sensitivity across the visual field.

To address this, two participants (C5 and C6) completed a modified version of the contrast sensitivity task in which they made eye movements either by following a dot moving randomly at a radius of 2° or 5° around fixation, or by self-initiated very large eye movements. Eye movements across these conditions (Figure 7, bottom row; Figure 8, bottom row), were quantified using BCEA (**C5** – Central fixation: mean±SD = 0.57±0.11 deg^2^, 2° eye motion: 2.69±0.48 deg^2^, 5° eye motion: 20.3±1.32 deg^2^, random eye motion: 133.7±23.36 deg^2^; **C6** – Central fixation: 0.96±0.56 deg^2^, 2° eye motion: 1.28±0.15 deg^2^). For reference, in severe (idiopathic) nystagmus, the eye movement variability along the vertical and horizontal planes is 1.08 deg and 1.60 deg, respectively (Tailor et al., 2021). Assuming a moderate correlation between axes (𝜌 = 0.3), the average fixation stability would equate to a BCEA of ∼21.46 deg^2^ (i.e., ∼5° eye motion condition in our data).

**Figure 7.**
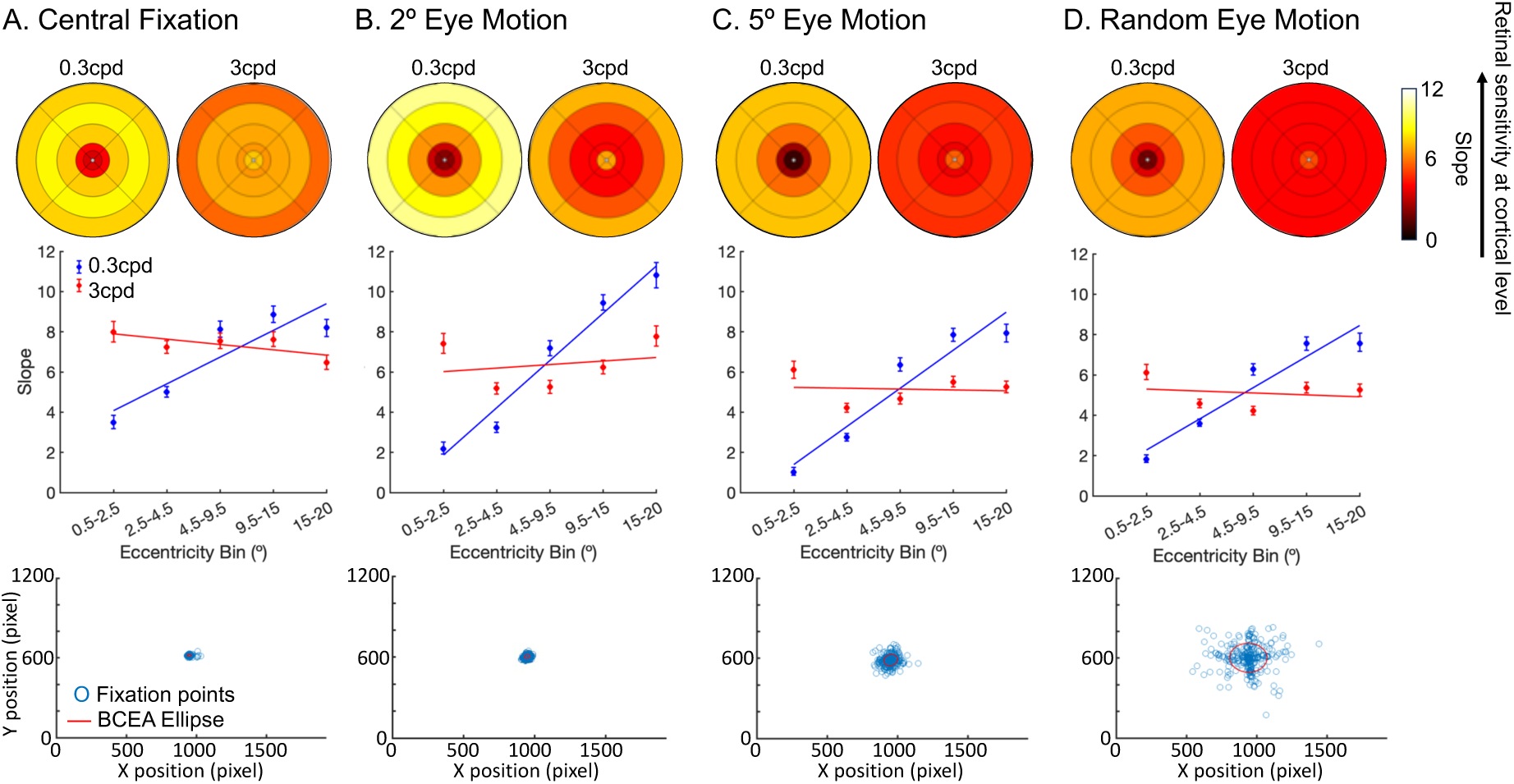
Effect of eye movements on slope estimation, in participant C5. Each column represents a distinct eye movement condition: A: central fixation, B: 2° eye motion, C: 5° eye motion, and D: random eye motion. pRF estimates were used to relate brain responses to visual field positions. Top row: Heatmaps of slope distributions for low (0.3 cycles per degree, cpd) and high (3 cpd) spatial frequency stimuli, with warmer colors indicating steeper slopes . Middle row: Slope values scattered across eccentricity bins (0.5–2.5°, 2.5–4.5°, 4.5–9.5°, 9.5–15°, and 15–20°), illustrating how cortical sensitivity varies across the visual field for low (blue) and high (red) spatial frequencies. Error bars represent 95% confidence interval. Bottom row: Fixation stability plots from one out of 3 runs, including fixation points (blue) and BCEA ellipses (red). BCEA values quantify gaze dispersion: Central Fixation (0.91 deg²), 2° eye motion (2.43 deg²), 5° eye motion (18.41 deg²), and random eye motion (130.79 deg²). Despite increased eye movements and reduced slope values – particularly for high spatial frequency stimuli – the overall pattern of cortical sensitivity remains consistent across conditions.

**Figure 8.**
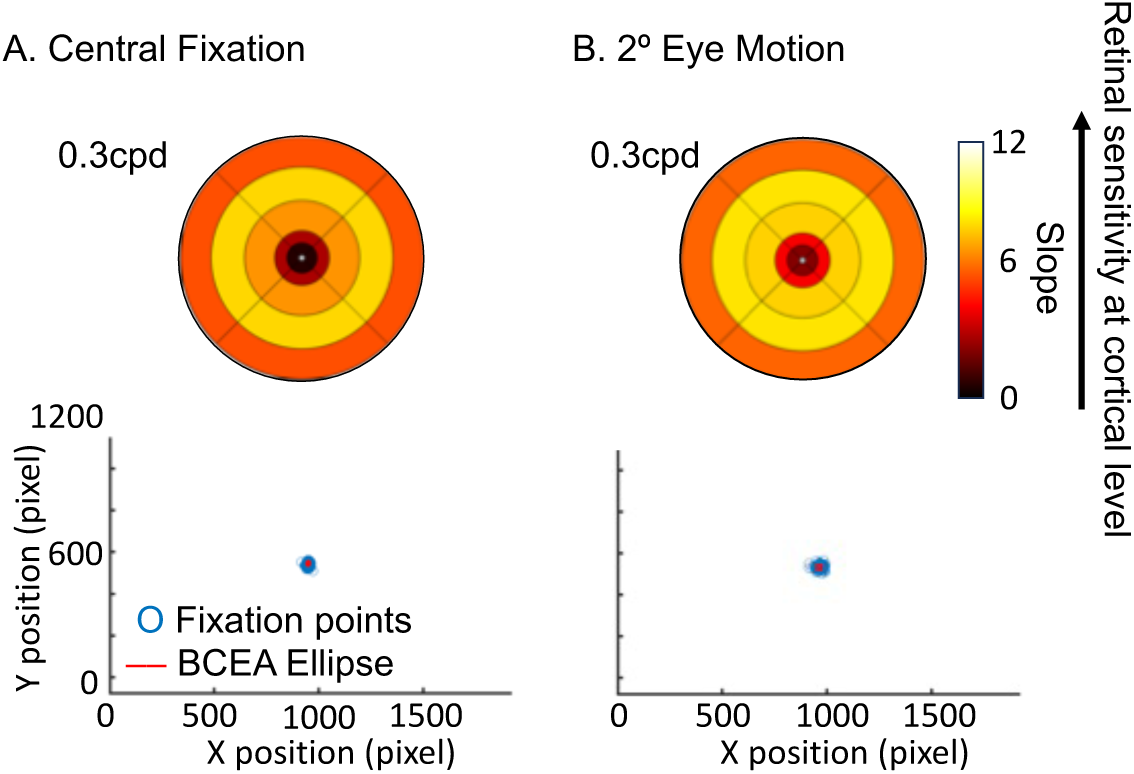
Effect of eye movements on slope estimation, in participant C6. Each column represents a distinct eye movement condition, A: central fixation. B: 2° eye motion. pRF estimates were used to relate brain responses to visual field positions. Top row: Heatmaps of slope distributions for low (0.3 cycles per degree, cpd) and high (3 cpd) spatial frequency stimuli, with warmer colors indicating steeper slopes. Bottom row: Fixation stability plots from one out of 3 runs, including fixation points (blue) and BCEA ellipses (red). BCEA values quantify gaze dispersion: Central Fixation (0.76 deg²) and 2° eye motion (1.23 deg^2^).

Despite these very large levels of eye movements, we observed that the overall cortical contrast sensitivity spatial pattern across eccentricity remained remarkably consistent (Figure 7, top and middle rows; Figure 8, top row). However, at the most extreme movements, contrast sensitivity estimates (slope values) were lower; and while the overall cortical visual field map structure was still clearly present for low spatial frequencies, it appeared more flattened for 3cpd, suggesting reduced sensitivity of our measure for large eye movement and high spatial frequency stimuli.

### A test-case of simulated loss of visual inputs

In the previous sections, we showed that the slope of a square root function provides a reliable measure of contrast sensitivity in the brain of healthy controls. But can this brain-level model also quantify loss of visual inputs? To test this, we first simulated an artificial scotoma in one normal sighted participant, by (a) masking a region of the visual field with a grey, annular ring, covering 3°-8° eccentricity (Figure 9A), and (b) masking the upper-right visual quadrant using a grey quarter-sector overlay (Figure 10A). We expect smaller slope values in V1 neuronal populations that would under normal circumstances encode that part of the visual space.

**Figure 9.**
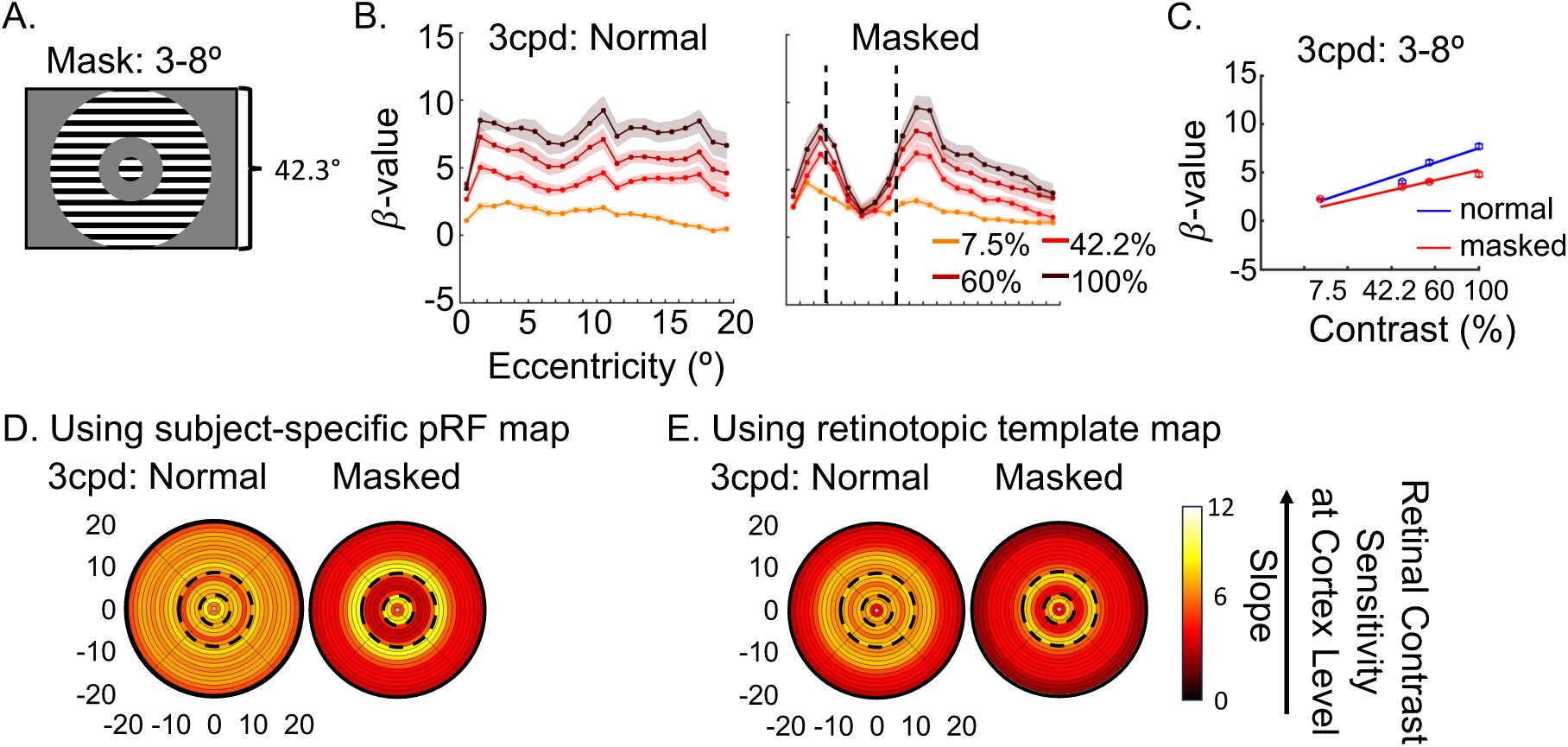
V1 sensitivity (i.e., slope) to 3cpd condition, in the region of the artificial scotoma (3-8°) in participant C5. A: Masked region between 3-8°, representing the artificial scotoma. B: Averaged 𝛽-values between 0.5°-20° eccentricities in V1, under normal and masked conditions. 20 eccentricity bins of 1° were selected based on pRF estimates. The color scale indicates the presented contrast levels, and the black dotted lines mark the boundaries of the artificial scotoma. Shaded area represents 95% confidence interval. C: Average 𝛽-values versus contrast levels in V1 neurons encoding the region of artificial scotoma (3°-8°), under normal (blue) and masked (red) conditions. Lines correspond to the model fitted to the data. Error bars (masked by dots) represent 95% confidence interval. D: Slopes projected back onto the visual field under normal and masked conditions. pRF estimates were used to relate brain responses to visual field locations and to create the 1° eccentricity bins. The boundaries of the simulated scotoma region are represented by dotted lines. The color scale indicates slope estimates, with higher values (warmer colors) corresponding to higher V1 contrast sensitivity. E: Slopes plotted back into the visual space using the calibrated Benson template instead of pRF estimates.

**Figure 10.**
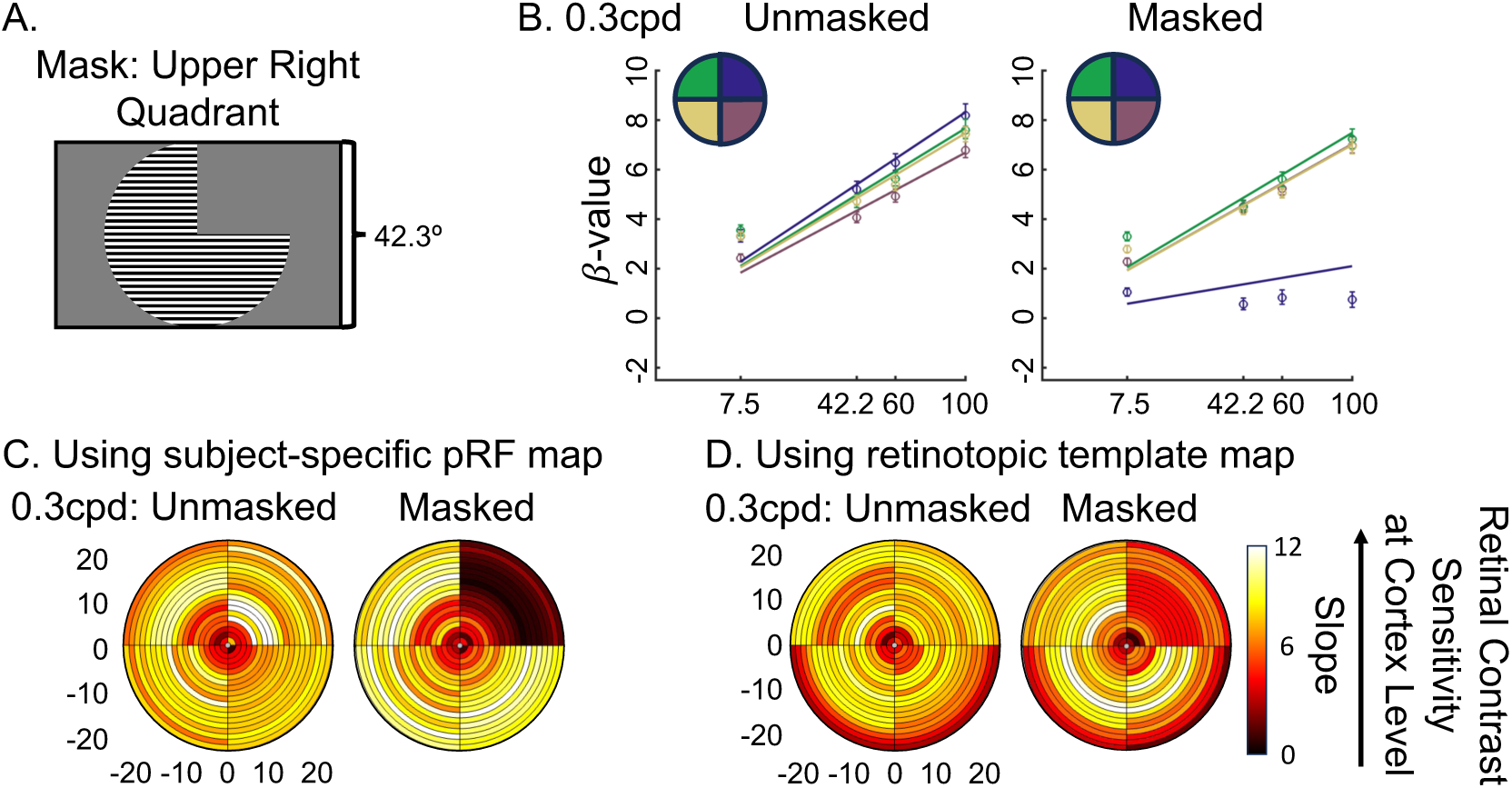
V1 sensitivity (i.e., slope) to 0.3cpd condition, in the region of the artificial scotoma (upper right quadrant) in participant C5. **A:** Masked upper right quadrant region, representing the artificial scotoma. **B:** Average 𝛽-values versus contrast levels in V1 neurons encoding each of the four quadrants: upper left (yellow), upper right (cyan), bottom left (red), and bottom right (blue). Lines correspond to the model fitted to the data. Error bars represent 95% confidence interval. **D:** Slopes projected back onto the visual field under normal and masked conditions. pRF estimates were used to relate brain responses to visual field locations and to create the 1° eccentricity bins. The color scale indicates slope estimates, with higher values (warmer colors) corresponding to higher V1 contrast sensitivity. **E:** Slopes plotted back into the visual space using the calibrated Benson template instead of pRF estimates.

As expected, we observed reduced responses in V1 locations corresponding to the artificial scotoma (Figures 9 and 10), with increased responses along the edges of the mask for the ring scotoma condition (Figure 9B). This artificial loss of visual input was also clearly present in the cortical contrast sensitivity estimate, with significantly reduced slope steepness in V1 between 3-8° for the ring scotoma condition (Figure 9C&D) and in the upper-right quadrant for the quarter-sector scotoma condition (Figure 10B&C). Additionally, we could recover this scotoma using the calibrated Benson template, although less accurately (Figures 9E and 10D). These results show that this measure of V1 contrast sensitivity is sensitive enough to detect loss of visual inputs in the brain at an individual level, when a complete local loss of sight is simulated, and that this approach does not crucially rely on pRF mapping data from the individual. This supports the utility of our approach in recovering patterns of vision loss and recovery at a cortical level.

### Case of severe central vision loss

We next tested if brain-based cortical contrast sensitivity could recover visual field loss in a clinical patient with Leber Hereditary Optic Neuropathy (LHON; M, 45.5 years old, mutation: m.11778G>A, disease duration: 36.1 years). LHON is a neurodegenerative disease caused by mitochondrial DNA mutations, leading to acute loss of retinal ganglion cells across large visual field areas. LHON patients often present with dense, central scotoma extending up to 10-15 degrees around fixation.

As vision loss often varies between eyes, the patient underwent monocular testing for the fMRI contrast sensitivity task (tested spatial frequencies: 0.3cpd and 1cpd). For comparison, a behavioral visual field perimetry map was obtained with the Compass fundus perimeter (CenterVue, Padova, Italy) for each eye, using a 24-2 testing grid with a Goldmann size III target (54 locations: 52 from the 24-2 grid, 1 at fixation, and 1 at the physiological blind spot). The Compass actively tracks the retina to compensate for fixation loss, automatically repositioning targets based on current eye position. Sensitivity thresholds at each location were determined using the ZEST projection strategy implemented in the device. The patient, who had some preserved central vision, could see the fixation targets in behavioral testing and the scanner, making them an ideal participant for methods evaluation.

The Compass perimetry map revealed drastic loss of sensitivity across the visual field (Figure 11 – top row). For the left eye, the left hemifield was more affected than the right, with most severe loss in the upper-left quadrant. In contrast, the right eye showed more reduced sensitivity in the right hemifield, particularly in the upper-right quadrant.

**Figure 11.**
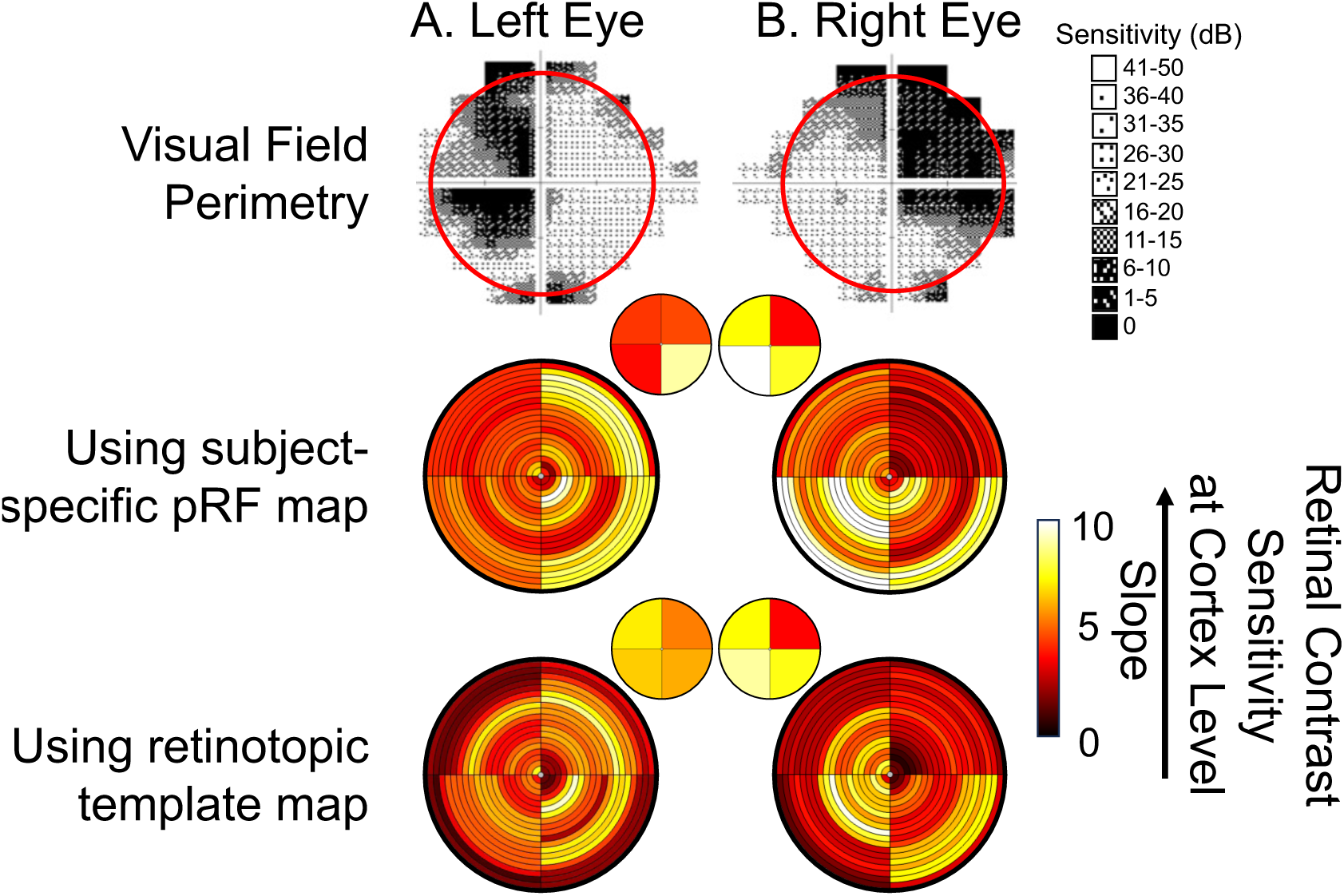
Correspondence between visual field perimetry map and V1 sensitivity map, in a patient with Leber Hereditary Optic Neuropathy. **A:** Left eye. **B:** Right eye. **Top row:** Gray scale visual field map obtained with the Compass fundus perimeter (CenterVue, Padova, Italy), using Standard Perimetry display convention. Sensitivity is represented with symbols and related dB intervals, with larger values describing better sensitivity. Red circle indicates the area of the visual space stimulated in the fMRI contrast sensitivity task. **Middle row:** Heatmap of V1 sensitivity (i.e., slopes) across the visual field. Here, we divided the space between the upper-left, upper-right, lower-left and lower-right quadrants (as opposed to the left, right, upper and lower quadrants used in previous sections), to match the layout of visual field perimetry maps. We also only show the responses to low spatial frequency condition (0.3cpd) for visualization. Slope values were averaged for each visual quadrant and 1° eccentricity bin and projected back into the visual space, using the subject-specific pRF map. Color scale corresponds to the steepness of the slope estimate. For better visualization of sensitivity differences across quadrants, we also averaged the slopes between 0.5-20° eccentricities in each quadrant, generating a single slope value for each visual quadrant (inset heat plots). The range of slope values were reduced on the color scale for these inset plots. **Bottom row:** Heatmap of V1 sensitivity (i.e., slopes) across the visual field in response to the 0.3cpd condition, using the Benson retinotopic template map instead of the pRF data.

We found correspondence between affected and non-affected visual field regions across the Compass and the fMRI contrast sensitivity measures (Figure 11). Here, V1 contrast sensitivity (i.e., slope values) was computed for the upper-left, upper-right, lower-left and lower-right quadrants to match the layout from the Compass visual field maps. Using the brain-based approach, we recovered sensitivity loss in the upper-right visual quadrant for the right eye at the cortical level, reflected in smaller cortical contrast sensitivity slopes in V1 neuronal populations encoding that area. For the left eye, cortical contrast sensitivity was also lower in the upper-left quadrant of the visual field, but differences were less pronounced. This is in line with the observation that the left eye was the better eye, with substantially less severe vision loss in the affected regions (lighter shading, Figure 11 – top row). When using the calibrated Benson atlas instead of pRF mapping data to link neuronal responses to visual locations, this pattern was retained for the right eye, but the more subtle pattern in the left eye became noisier.

By combining large-field stimulation and contrast sensitivity modeling in fMRI, with the cortical magnification-adjusted Benson atlas, these data show how brain-based measures could provide a tool to characterize vision loss in patients without need for demanding tasks or sustained precise fixation, while highlighting challenges of detecting more subtle vision loss and pinpointing it to visual locations.

## Discussion

In this study, we aimed to develop and validate an fMRI-based method for measuring cortical contrast sensitivity across a broad extent of the visual field in V1, addressing the limitations of traditional visual field tests and existing neuroimaging approaches. To achieve this, we implemented a large-field presentation set-up in fMRI, allowing us to stimulate the visual cortex up to 20° eccentricity. By integrating population receptive field (pRF) modeling with contrast sensitivity measurements of the BOLD response to large-field sinusoidal gratings varying in contrast levels and spatial frequencies, we created a quantitative, sensitive, and reliable measure of cortical contrast sensitivity. Our key findings demonstrate that this approach can accurately quantify known variations in V1 contrast sensitivity across eccentricity and polar angle, recover simulated and pathological visual field losses, and produce consistent and reliable measures both at the group and individual participant levels. Crucially, the ability to visualize these sensitivity gradients was made possible by the large peripheral coverage provided by our large-field stimulation set-up. Such coverage is particularly important for clinical applications, as it enables the detection of visual field losses beyond the macula (i.e., beyond 10° eccentricity) and the evaluation of residual peripheral vision in patients with macular-restricted damage. In doing so, this work provides a useful tool for advancing both basic visual neuroscience and translational research in clinical populations.

### Validation of Sensitivity Variations Across the Visual Field

Our method successfully captured the expected variations in cortical contrast sensitivity across the visual field. Consistent with previous behavioral and fMRI studies (Aghajari et al., 2020; Broderick et al., 2022; D’Souza et al., 2016; Henriksson et al., 2008; Rovamo et al., 1984; Rovamo & Virsu, 1979; Welbourne et al., 2018), we observed that V1 neuronal populations encoding the central visual field exhibited higher contrast sensitivity for high spatial frequency (SF) stimuli, while those encoding peripheral regions were more sensitive to low SF stimuli. This aligns with the understanding that spatial frequency tuning varies across the visual field due to smaller pRF sizes and larger cortical areas dedicated to central vision (Himmelberg et al., 2023).

Additionally, our study identified anisotropies in contrast sensitivity across visual field quadrants, mirroring findings from previous behavioral studies (Barbot et al., 2021; Carrasco et al., 2022; Himmelberg et al., 2020). Specifically, we found enhanced cortical sensitivity in V1 neuronal populations responding to horizontal versus vertical locations and to lower versus upper visual field locations. Unlike earlier fMRI studies (Liu et al., 2006; O’Connell et al., 2016), which either could not assess horizontal-vertical effects or used smaller region selections, our use of large-field stimuli enabled the detection of both upper-lower and horizontal-vertical asymmetries. Moreover, by modeling cortical contrast sensitivity across multiple contrast levels, we were able to identify subtle variations in V1 neuron sensitivity that single-threshold approaches might overlook.

We also observed intriguing variability in cortical visual field maps across healthy controls, and this variability was consistent across measures. This may reflect genuine individual differences in visual sensitivity that are relevant for behavioral performance. Alternatively, it could arise from factors such as local signal-to-noise differences driven by anatomical variability. However, the fact that maps derived from different spatial stimulus conditions showed markedly different patterns argues against a purely anatomical explanation and suggests that at least part of the variability is functional. Despite this inter-subject variability, variations in cortical contrast sensitivity across eccentricities and visual field quadrants were significant at the individual level indicating high sensitivity.

### Reliability and Sensitivity Across Sessions and Robustness to Eye Movements

We also showed that our approach was reliable over time. We found consistent activation patterns and moderate-to-high correlations in slope estimates for the 0.3cpd condition across two sessions, significantly higher than correlations across different stimuli conditions (i.e., 0.3cpd versus 3cpd). High sensitivity and reliability are essential for detecting longitudinal changes in clinical groups, such as those occurring before and after sight-rescuing interventions. Fitting a contrast response function to the BOLD signal might offer robust estimates of visual function by minimizing the impacts of overall BOLD amplitude variability and arbitrary thresholds.

Crucially, one advantage of cortical visual field mapping is that the maps are inherently centered on the foveal confluence, providing a stable reference point for comparing responses across eccentricities. When combined with large-field, spatially homogeneous stimuli, this anchoring means that our approach should remain robust to moderate fixation variability and still quantify sensitivity changes across the visual field – provided that fixation instability does not exceed the stimulus extent (40° diameter).

When measuring the impact of eye movements, we found that spatial sensitivity patterns were largely preserved, even for extreme eye movements (emulating severe nystagmus). However, under the most extreme conditions, sensitivity estimates (i.e., slope values) were reduced, especially for high spatial frequency (SF) stimuli. This likely reflects image blurring from large rapid eye movements, which degrades high-SF inputs and shifts activation toward neurons tuned to lower SFs. This aligns with evidence that nystagmus and large saccades impair perception of fine detail and grating stimuli due to retinal image slip (Abadi & Bjerre, 2002; Dickinson & Abadi, 1985; Hertle et al., 2017; Randall et al., 2020). While classic findings report suppression of low-SF signals during saccades (Burr et al., 1994; Ross et al., 2001), our results suggest that high SF sensitivity may be more vulnerable to large eye movements when participants are presented with 2Hz phase-flickering gratings. Further validation in clinical groups with naturally-occurring fixation instability would further strengthen these conclusions.

### Mapping Simulated and Pathology-Driven Vision Loss

Our method successfully identified both simulated retinal loss in a healthy volunteer and real visual field loss in a patient with Leber Hereditary Optic Neuropathy (LHON). The signal drop observed in response to masking portions of the visual field in the healthy control was both large and significant at the individual level, as demonstrated by non-overlapping 95% confidence intervals (Figures 9B-C and 10B). This provides proof-of-concept evidence that our approach can detect signal changes in individual patients, which is a critical requirement for clinical translation.

Unlike previous fMRI studies that used high-contrast stimuli (Farahbakhsh et al., 2022; Pawloff et al., 2023; Ritter et al., 2019), which may not accurately represent partial vision loss due to potential saturation effects and the stimulation of less sensitive retinal cells, our use of multiple contrast levels offers a more nuanced assessment of cortical contrast sensitivity. Combined with the large-field set-up allowing stimulation up to 20° eccentricity, this approach may be particularly well-suited for evaluating treatment efficacy in cases of widespread and variable vision loss.

Future work will focus on further validating reconstruction accuracy under controlled conditions, including simulated scotomas of varying severity and location, expanding testing to larger patient cohorts, and establishing a normative dataset to contextualize patient data.

### Limitations of pRF Mapping, Advances in Structure-Based Templates, and Future Directions in Brain-Based Visual Field Mapping

One limitation of this brain-based approach to visual field testing, is its reliance on pRF mapping, which requires precise fixation and can be challenging to acquire in visually impaired patients. To mitigate this, we investigated the use of retinotopic templates based on anatomical landmarks (Benson et al., 2014; Benson & Winawer, 2018), thereby reducing the need for pRF data collection and facilitating the mapping of brain responses to visual field positions. Particularly at large eccentricities however, we initially observed inaccuracies between the template and individual retinotopy eccentricity estimates which led to substantial distortions in cortical visual field maps due to cortical magnification (see Figure A4 in *Appendix* section). To address this, we adjusted the Benson eccentricity estimates to align with the cortical magnification scaling function (Horton & Hoyt, 1991).

With this structure-based atlas, we successfully replicated key variations in visual field function (across eccentricity and polar quadrants), although sensitivity to more subtle differences (e.g., upper versus lower quadrant anisotropy) was reduced. This reduction may partly stem from differences in ROI definitions: a manually delineated V1 mask was used for the pRF-based data, while the Benson atlas mask was used for the adjusted Benson atlas analysis. Such differences could introduce minor error beyond the atlas/pRF mapping itself due to differences in the vertices included by each mask.

Beyond ROI considerations, we still observed differences in cortical sensitivity between pRF mapping and the adjusted Benson atlas – particularly in the periphery. Several factors likely contribute to this. First, individual differences in the relationship between cortical structure and retinotopy are not fully captured by the template. Second, the Benson atlas has never been fit with empirical data more eccentric than approximately 20°, which naturally limits its precision in the far periphery. Third, because of cortical magnification, any small inaccuracy at larger eccentricities has a disproportionately large effect, making peripheral vertices more susceptible to mis-assignment than central ones. These influences introduce systematic distortions in cortical visual field maps rather than random noise and thus remain consistent across timepoints – an important point when assessing longitudinal changes (e.g., ageing or gene-therapy interventions). Importantly, the spatial gradients in cortical contrast sensitivity were preserved across both the pRF and Benson atlas approaches, indicating that minor ROI differences do not affect our conclusions. Together, these findings show that the Benson atlas remains a useful alternative when pRF mapping is not feasible.

It is also promising that cortical structure-based templates can successfully recover visual field loss through simulation or the neurodegenerative disease LHON. Critically, in patient groups with severe impairment such as LHON, where it is often very challenging or impossible to obtain reliable measures, even imperfect mapping between the brain and visual field could offer important improvements over available solutions for tracking functional change, and where they stem from. Further enhancing the alignment between retinotopic template atlases and individual retinotopic tuning could improve this approach further, for example, by integrating them with functional measures using Bayesian methods (Benson & Winawer, 2018). In parallel, geometric deep learning frameworks such as DeepRetinotopy (Ribeiro et al., 2021) could also offer anatomy-driven predictions from structural MRI, and combining these strategies may yield more accurate and generalizable retinotopic reconstructions.

While our method demonstrated high accuracy and reliability in sighted controls, further validation in patients is necessary to ensure the generalizability of our approach to measuring pathology. Next steps in this work will therefore involve testing larger patient cohorts with diverse forms of vision loss, validating the approach for tracking pathology over time, and investigating how cortex-based visual field measures relate to and complement other visual field and retinal integrity indices including Compass measures and OCT-derived retinal layer thickness. For example, while we reasoned that using a single high contrast stimulus is likely to give less reliable measures for various reasons outlined above, this needs to be tested empirically. It also remains unclear whether the square root contrast response function is optimal for patients, who are likely to show different non-linearities in their contrast response. In addition, comparing the cortical contrast sensitivity modeling approach presented here with alternative framework (e.g., neural contrast sensitivity functions proposed by Roelofzen et al., 2025) will be another interesting direction for future research. Finally, more extensive mapping of the spatial frequency and contrast sensitivity spaces could offer additional informative measures such as cortical acuity and spatial frequency preferences, provided such measures are feasible with patient-friendly protocols.

## Conclusion

In sum, brain-based contrast sensitivity measures could significantly enhance the accurate characterization of visual field function in blinding diseases, by addressing limitations of traditional visual field tests that are subjective, time-consuming, and challenging for patients with severe sight loss. By providing a reliable, quantitative measure of cortical contrast sensitivity across the visual field, fMRI may help facilitate the monitoring of disease progression and recovery, including interventions like gene therapy. Combined with our large-field stimulation setup and fixation free approaches to linking cortical sensitivity estimates to visual field locations, this method is well-suited for a wide range of visual impairments, addressing a pressing challenge in the evaluation of current clinical trials. Additionally, linking brain-based variations in function across the visual field to behavioral performance (e.g., perimetry, microperimetry) and retinal structure (fundus imaging, retinal thickness from Optical Coherence Tomography), could help bridge the gap between neural measures and functional outcomes. Such integration would provide deeper insights into developmental, learning, and vision loss mechanisms.

In conclusion, integrating cortical contrast sensitivity testing with large-field stimulation and a calibrated retinotopic atlas provides a promising tool for characterizing visual field impairments, and could offer a valuable addition to existing measures of visual field loss.

## Acknowledgements

This work was funded by Moorfields Eye Charity PhD Studentship GR001315 (London, UK), the Wellcome Career Development Award (306332/Z/23/Z; London, UK), and the Birkbeck-UCL Centre for NeuroImaging (BUCNI; London, UK). PYWM is supported by an Advanced Fellowship Award (NIHR301696) from the UK National Institute of Health Research (NIHR). PYWM also receives funding from Fight for Sight (UK), the Isaac Newton Trust (UK), Moorfields Eye Charity (GR001376), the Addenbrooke’s Charitable Trust, the National Eye Research Centre (UK), the International Foundation for Optic Nerve Disease (IFOND), the NIHR as part of the Rare Diseases Translational Research Collaboration, the NIHR Cambridge Biomedical Research Centre (NIHR203312), and the NIHR Biomedical Research Centre based at Moorfields Eye Hospital NHS Foundation Trust and UCL Institute of Ophthalmology (NIHR203322). The views expressed are those of the author(s) and not necessarily those of the NHS, the NIHR or the Department of Health. We extend our thanks to Simon Richardson and Oliver Josephs for their assistance in the design and conception of the large-field set-up.

## Data Availability Statement

All post-processed and anonymised data are available on Zenodo (https://doi.org/10.5281/zenodo.19051439).

Codes to reproduce the figures from the provided dataset are available on GitHub (https://github.com/HugoCWB/ChowWingBomEtAl_2025_eLife).

Codes to adjust the Benson maps eccentricity is available on GitHub (https://github.com/HugoCWB/AdjustEccTool). A Docker image for the Benson eccentricity adjustment toolbox is also available on Zenodo (https://doi.org/10.5281/zenodo.18770065).

For any queries or raw data requests, please email the corresponding author.

## Appendix

### A. Large-Field set-up

**Figure A1.**
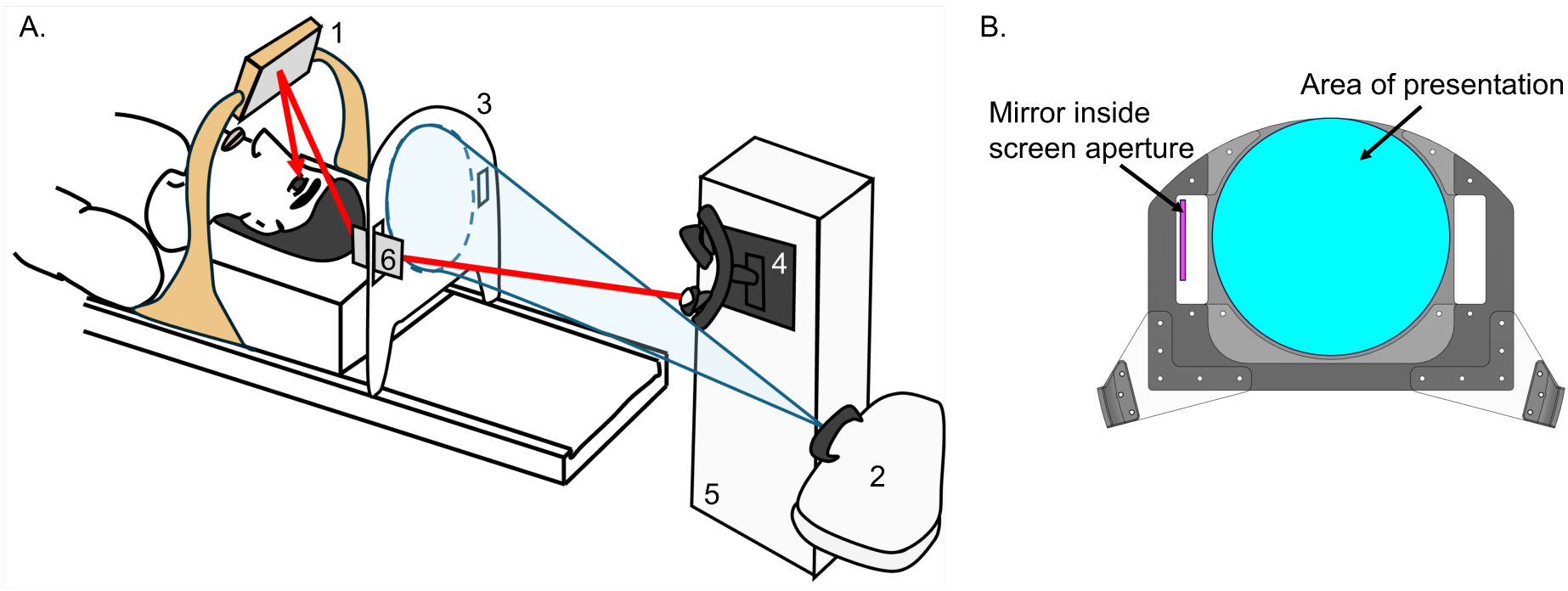
Schematic of Large-Field Set-Up. A: The participant lies on the scanner bed in the 64-channel coil without top to reduce obstruction and views the 40° screen via a mirror above their face (1). Stimuli are displayed using back-projection from a projector (2) outside the room, onto a screen (3) positioned behind the participant’s head. The top of the screen follows the scanner bore curvature to maximize field of view. Monocular eye-tracking is achieved by mounting the illuminator and camera of the eye-tracker (4) vertically on a support (5) at the back of the scanner bore. An image of the eye is obtained via a dual-mirror set-up, including a small mirror inside an aperture cut on the side of the screen (6) and the participant mirror (1). If the right eye is being tracked, the eye-tracker is placed on the left side of the scanner bore, and the eye-tracker mirror (6) is placed in the right screen aperture. B: Schematic of the large-field screen, with the area of stimulus presentation shown in cyan and the mirror used for eye-tracking in magenta. The mirror is positioned within one of the two rectangular apertures cut on either side of the screen.

#### B. Individual Level

To test for contrast sensitivity differences across the visual field in individual participants, we also employed a multilevel modeling approach using the lme4 package in R (Bates et al., 2015; R Core Team, 2023). This approach reduces the impact of vertex-level variability and redundancy. The model included the square root of stimulus contrast as a continuous fixed-effect predictor, and eccentricity, polar location, and spatial frequency as categorical fixed-effect predictors, along with all possible interactions. Again, the intercept was fixed at 0. The model also included random slopes for the square root contrast and the interaction between square root contrast and voxel-specific factors and was fit separately for each participant. In the formula notation used in the R package lme4, the model would be defined as:

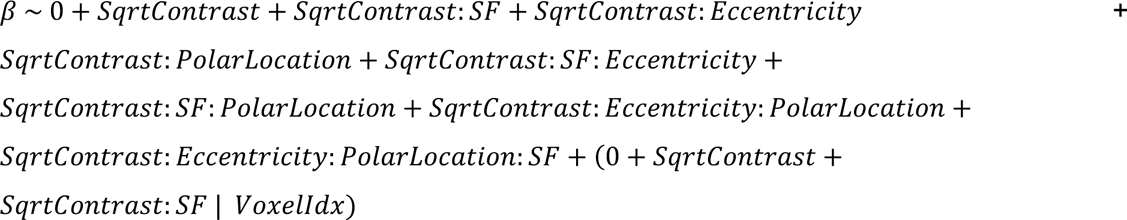

**Table A1.** Results from the multilevel modelling approach in each individual participant. pRF mapping was used to link brain responses to visual field locations.

| SubID | Effects | Df | Sum Sq | Mean Sq | F-value | Pr(>F) | Signif. |
| --- | --- | --- | --- | --- | --- | --- | --- |
| C1 | SF | 1 | 3720 | 3720 | 72.49 | < 2e-16 | *** |
|  | Eccentricity | 1 | 35003 | 35003 | 681.995 | < 2e-16 | *** |
|  | PolarLocation | 3 | 11258 | 3753 | 73.115 | < 2e-16 | *** |
|  | SF:Eccentricity | 1 | 23914 | 23914 | 465.938 | < 2e-16 | *** |
|  | SF:PolarLocation | 3 | 1546 | 515 | 10.041 | 1.32E-06 | *** |
|  | Eccentricity:PolarLocation | 3 | 200 | 67 | 1.297 | 0.274 |  |
|  | SF:Eccentricity:PolarLocation | 3 | 1158 | 386 | 7.519 | 5.04E-05 | *** |
|  | Residuals | 12378 | 635290 | 51 |  |  |  |
| C2 | SF | 1 | 23704 | 23704 | 743.628 | < 2e-16 | *** |
|  | Eccentricity | 1 | 13503 | 13503 | 423.609 | < 2e-16 | *** |
|  | PolarLocation | 3 | 46442 | 15481 | 485.647 | < 2e-16 | *** |
|  | SF:Eccentricity | 1 | 10587 | 10587 | 332.135 | < 2e-16 | *** |
|  | SF:PolarLocation | 3 | 2981 | 994 | 31.169 | < 2e-16 | *** |
|  | Eccentricity:PolarLocation | 3 | 2881 | 960 | 30.126 | < 2e-16 | *** |
|  | SF:Eccentricity:PolarLocation | 3 | 373 | 124 | 3.899 | 8.52E-03 | ** |
|  | Residuals | 11604 | 369897 | 32 |  |  |  |
| C3 | SF | 1 | 27 | 27 | 1.374 | 0.2411 |  |
|  | Eccentricity | 1 | 578 | 578 | 29.019 | 7.33E-08 | *** |
|  | PolarLocation | 3 | 7315 | 2438 | 122.52 | < 2e-16 | *** |
|  | SF:Eccentricity | 1 | 14719 | 14719 | 739.59 | < 2e-16 | *** |
|  | SF:PolarLocation | 3 | 257 | 86 | 4.311 | 4.80E-03 | ** |
|  | Eccentricity:PolarLocation | 3 | 2225 | 742 | 37.261 | < 2e-16 | *** |
|  | SF:Eccentricity:PolarLocation | 3 | 510 | 170 | 8.543 | 1.16E-05 | *** |
|  | Residuals | 9860 | 196225 | 20 |  |  |  |
| C4 | SF | 1 | 25290 | 25290 | 521.73 | < 2e-16 | *** |
|  | Eccentricity | 1 | 3970 | 3970 | 81.89 | < 2e-16 | *** |
|  | PolarLocation | 3 | 34475 | 11492 | 237.08 | < 2e-16 | *** |
|  | SF:Eccentricity | 1 | 75054 | 75054 | 1548.38 | < 2e-16 | *** |
|  | SF:PolarLocation | 3 | 2514 | 838 | 17.29 | 3.40E-11 | *** |
|  | Eccentricity:PolarLocation | 3 | 16417 | 5472 | 112.89 | < 2e-16 | *** |
|  | SF:Eccentricity:PolarLocation | 3 | 3921 | 1307 | 26.97 | < 2e-16 | *** |
|  | Residuals | 9896 | 479688 | 48 |  |  |  |
| C5 | SF | 1 | 554 | 554 | 16.332 | 5.37E-05 | *** |
|  | Eccentricity | 1 | 6717 | 6717 | 198.161 | < 2e-16 | *** |
|  | PolarLocation | 3 | 886 | 295 | 8.711 | 9.12E-06 | *** |
|  | SF:Eccentricity | 1 | 13436 | 13436 | 396.384 | < 2e-16 | *** |
|  | SF:PolarLocation | 3 | 4753 | 1584 | 46.744 | < 2e-16 | *** |
|  | Eccentricity:PolarLocation | 3 | 3023 | 1008 | 29.727 | < 2e-16 | *** |
|  | SF:Eccentricity:PolarLocation | 3 | 2638 | 879 | 25.939 | < 2e-16 | *** |
|  | Residuals | 7728 | 261956 | 34 |  |  |  |
| C6 | SF | 1 | 6008 | 6008 | 246.099 | < 2e-16 | *** |
|  | Eccentricity | 1 | 21371 | 21371 | 875.448 | < 2e-16 | *** |
|  | PolarLocation | 3 | 11850 | 3950 | 161.806 | < 2e-16 | *** |
|  | SF:Eccentricity | 1 | 12327 | 12327 | 504.98 | < 2e-16 | *** |
|  | SF:PolarLocation | 3 | 714 | 238 | 9.745 | 2.04E-06 | *** |
|  | Eccentricity:PolarLocation | 3 | 1569 | 523 | 21.425 | 7.97E-14 | *** |
|  | SF:Eccentricity:PolarLocation | 3 | 762 | 254 | 10.409 | 7.80E-07 | *** |
|  | Residuals | 9360 | 228493 | 24 |  |  |  |
| C7 | SF | 1 | 10703 | 10703 | 278.799 | < 2e-16 | *** |
|  | Eccentricity | 1 | 19431 | 19431 | 506.149 | < 2e-16 | *** |
|  | PolarLocation | 3 | 18754 | 6251 | 162.84 | < 2e-16 | *** |
|  | SF:Eccentricity | 1 | 19611 | 19611 | 510.822 | < 2e-16 | *** |
|  | SF:PolarLocation | 3 | 153 | 51 | 1.325 | 2.64E-01 |  |
|  | Eccentricity:PolarLocation | 3 | 12080 | 4027 | 104.885 | < 2e-16 | *** |
|  | SF:Eccentricity:PolarLocation | 3 | 1380 | 460 | 11.983 | 7.90E-08 | *** |
|  | Residuals | 10794 | 414384 | 38 |  |  |  |
| Signif. codes: 0 '***' 0.001 '**' 0.01 '*' 0.05 '.' 0.1 ' ' 1 |  |  |  |  |  |  |  |

**Table A2.** Post-Hoc test on ANOVA visual quadrant position effect (PolarLocation) at the individual level. pRF mapping was used to relate brain responses to visual field locations.

| SubID | Effects | contrast | estimate | SE | df | t.ratio | p.value |  |
| --- | --- | --- | --- | --- | --- | --- | --- | --- |
| C1 | PolarLocation | HM>VM | 5.5138156 | 0.3278444 | 12354 | 16.818391 | 8.917786e-63 | *** |
|  |  | LVM>UVM | 0.5999187 | 0.2796053 | 12354 | 2.145591 | 0.03192504 | * |
|  |  | LHM>RHM | 0.7058217 | 0.1711807 | 12354 | 4.123254 | 3.759997e-05 | *** |
| C2 | PolarLocation | HM>VM | 5.533751 | 0.2284595 | 11580 | 24.22203 | 1.771783e-126 | *** |
|  |  | LVM>UVM | 4.387544 | 0.1852974 | 11580 | 23.67839 | 4.420105e-121 | *** |
|  |  | LHM>RHM | 2.288693 | 0.1336361 | 11580 | 17.12630 | 5.942464e-65 | *** |
| C3 | PolarLocation | HM>VM | 3.568024 | 0.2172194 | 9836 | 16.425896 | 7.786414e-60 | *** |
|  |  | LVM>UVM | 2.317875 | 0.1801032 | 9836 | 12.869704 | 1.339442e-37 | *** |
|  |  | LHM>RHM | 0.159411 | 0.1214377 | 9836 | 1.312697 | 0.1893155 |  |
| C4 | PolarLocation | HM>VM | 7.3408010 | 0.3070954 | 9872 | 23.903972 | 8.223761e-123 | *** |
|  |  | LVM>UVM | 3.1514598 | 0.2492052 | 9872 | 12.646043 | 2.250910e-36 | *** |
|  |  | LHM>RHM | -0.9417519 | 0.1794558 | 9872 | -5.247821 | 1.571031e-07 | *** |
| C5 | PolarLocation | HM>VM | 2.2997460 | 0.2874033 | 7704 | 8.001807 | 1.405261e-15 | *** |
|  |  | LVM>UVM | 0.2736311 | 0.2296279 | 7704 | 1.191628 | 0.2334437 |  |
|  |  | LHM>RHM | -0.2201730 | 0.1728344 | 7704 | -1.273896 | 0.2027387 |  |
| C6 | PolarLocation | HM>VM | 1.8040008 | 0.2259007 | 9336 | 7.985813 | 1.561107e-15 | *** |
|  |  | LVM>UVM | 3.2006411 | 0.1818476 | 9336 | 17.600678 | 3.057311e-68 | *** |
|  |  | LHM>RHM | 0.3046082 | 0.1340245 | 9336 | 2.272779 | 0.02306211 | * |
| C7 | PolarLocation | HM>VM | 5.9085493 | 0.3067623 | 10770 | 19.261005 | 2.637528e-81 | *** |
|  |  | LVM>UVM | 2.0334164 | 0.2617710 | 10770 | 7.767922 | 8.703410e-15 | *** |
|  |  | LHM>RHM | -0.3370417 | 0.1599345 | 10770 | -2.107373 | 0.03510831 | * |
| Signif. codes: 0 '***' 0.001 '**' 0.01 '*' 0.05 '.' 0.1 ' ' 1 |  |  |  |  |  |  |  |  |

#### C. Calibrated Benson Template

**Table A3.** Group-Level ANOVA analysis on slope estimates, as function of spatial frequency, eccentricities, and visual quadrant positions. The Benson template retinotopic map was used to relate brain responses to visual field locations.

| Measure of Interest: Slope | num Df | den Df | MSE | F | ges | Pr(>F) |  |
| --- | --- | --- | --- | --- | --- | --- | --- |
| SF | 1.0000 | 6.0000 | 31.436 | 0.6117 | 0.010887 | 0.4639018 |  |
| ECC | 2.1650 | 12.9901 | 32.694 | 4.7020 | 0.160027 | 0.0270342 | * |
| PolarLocation | 1.7729 | 10.6374 | 27.316 | 7.0357 | 0.163209 | 0.0129521 | * |
| SF:Eccentricity | 2.2706 | 13.6238 | 7.438 | 29.4822 | 0.221807 | 7.552e-06 | *** |
| SF:PolarLocation | 1.5885 | 9.5309 | 5.173 | 2.1025 | 0.009793 | 0.1783533 |  |
| Eccentricity: PolarLocation | 12.0000 | 72.0000 | 5.253 | 2.9645 | 0.096634 | 0.0020990 | ** |
| SF: Eccentricity: PolarLocation | 12.0000 | 72.0000 | 1.195 | 3.5212 | 0.028091 | 0.0003995 | *** |
| Signif. codes: 0 '***' 0.001 '**' 0.01 '*' 0.05 '.' 0.1 ' ' 1 |  |  |  |  |  |  |  |

**Table A4.** Post-hoc t-test analysis for the visual quadrant position effect (PolarLocation) and the interaction between spatial frequency and eccentricities (SF:Eccentricity) reported in Table A3. The Benson template retinotopic map was used to relate brain responses to visual field locations.

| Measure of Interest: Slope | contrast | estimate | SE | df | t.ratio | p.value |
| --- | --- | --- | --- | --- | --- | --- |
| PolarLocation | Horizontal>Vertical | 4.3378240 | 1.2101752 | 6 | 3.5844596 | 0.01158079 |
|  | Lower>Upper | 0.5249520 | 0.7053736 | 6 | 0.7442184 | 0.48485944 |
|  | Left>Right | -0.2265194 | 0.3922638 | 6 | -0.5774671 | 0.58462631 |
| SF:Eccentricity | 3cpd-0.3cpd: 0.5-2.5° | 4.5917619 | 0.7960846 | 6 | 5.7679320 | 0.001185113 |
|  | 3cpd-0.3cpd: 2.5-4.5° | 2.5801243 | 1.0192745 | 6 | 2.5313340 | 0.044599322 |
|  | 3cpd-0.3cpd: 4.5-9.5° | -0.2927635 | 0.9981812 | 6 | -0.2932969 | 0.779175269 |
|  | 3cpd-0.3cpd: 9.5-15° | -1.9782789 | 0.6042727 | 6 | -3.2738182 | 0.016953076 |
|  | 3cpd-0.3cpd: 15-20° | -2.2803080 | 0.6462658 | 6 | -3.5284368 | 0.012391168 |

**Figure A2.**
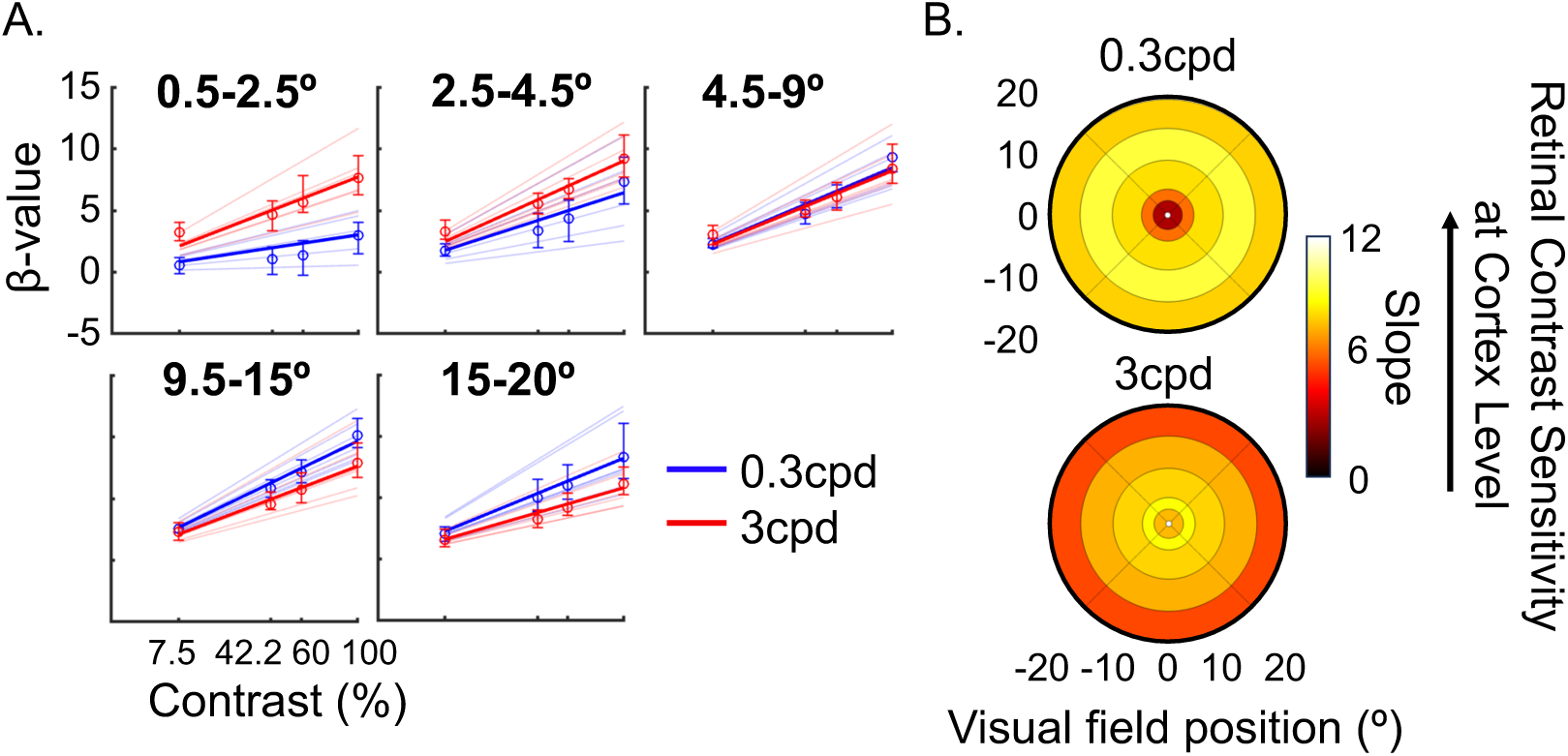
V1 contrast sensitivity (i.e., slopes) across 5 eccentricity bins (0.5°-2.5°, 2.5°-4.5°, 4.5°-9.5°, 9.5°-15°, and 15-20°), defined using the calibrated Benson atlas. **A:** β-values versus contrast levels for each eccentricity bin. Blue and red lines represent the contrast sensitivity model fits for the 0.3cpd and 3cpd conditions, respectively. Thinner lines correspond to individual fits. **B:** V1 contrast sensitivity index (i.e., slope) projected back into the visual space using the calibrated Benson atlas, for each eccentricity bin and spatial frequency condition. The color scale indicates slope estimates, with higher values representing higher V1 contrast sensitivity.

**Figure A3.**
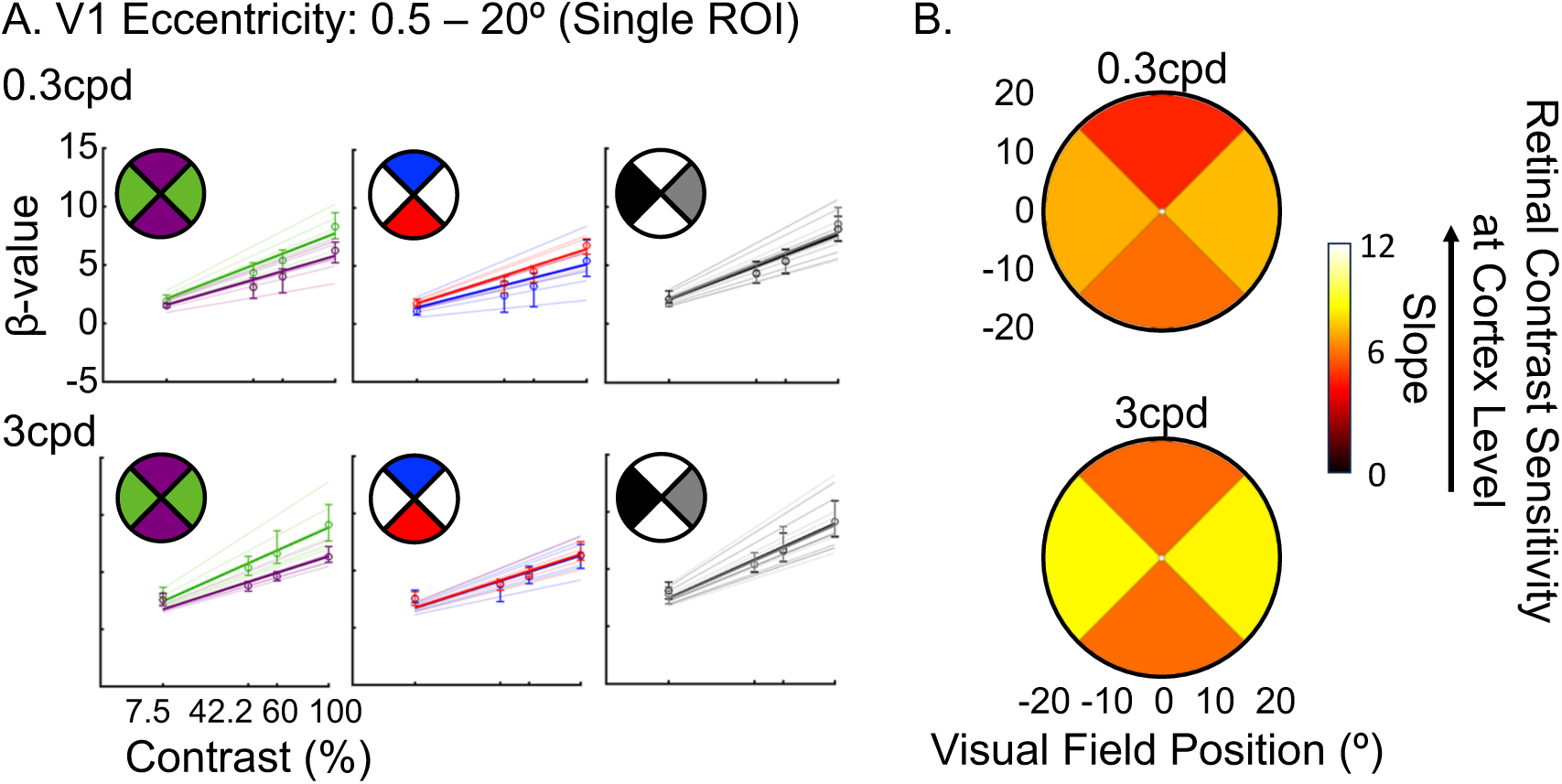
V1 contrast sensitivity (i.e., slopes) across visual field quadrants using the calibrated Benson atlas, for the 0.3cpd and 3cpd conditions. Visual quadrants were defined as ±45° regions around the cardinal meridians and slope values were selected for eccentricities between 0.5-20°. Horizontal quadrants are left and right, whilst upper and lower quadrants define the vertical quadrants. **A:** β-values versus contrast levels for each anisotropy effects: horizontal versus vertical, upper versus lower, and left versus right. Group-level model fits are in thick lines, whilst thinner lines correspond to individual fits. Error bars represent the 95% confidence intervals. **B:** V1 contrast sensitivity index (i.e., slope) projected back into the upper, lower, left and right visual field quadrants for the 0.3cpd and 3cpd conditions. The color scale represents slope estimates, with higher values indicating higher V1 contrast sensitivity.

**Figure A4.**
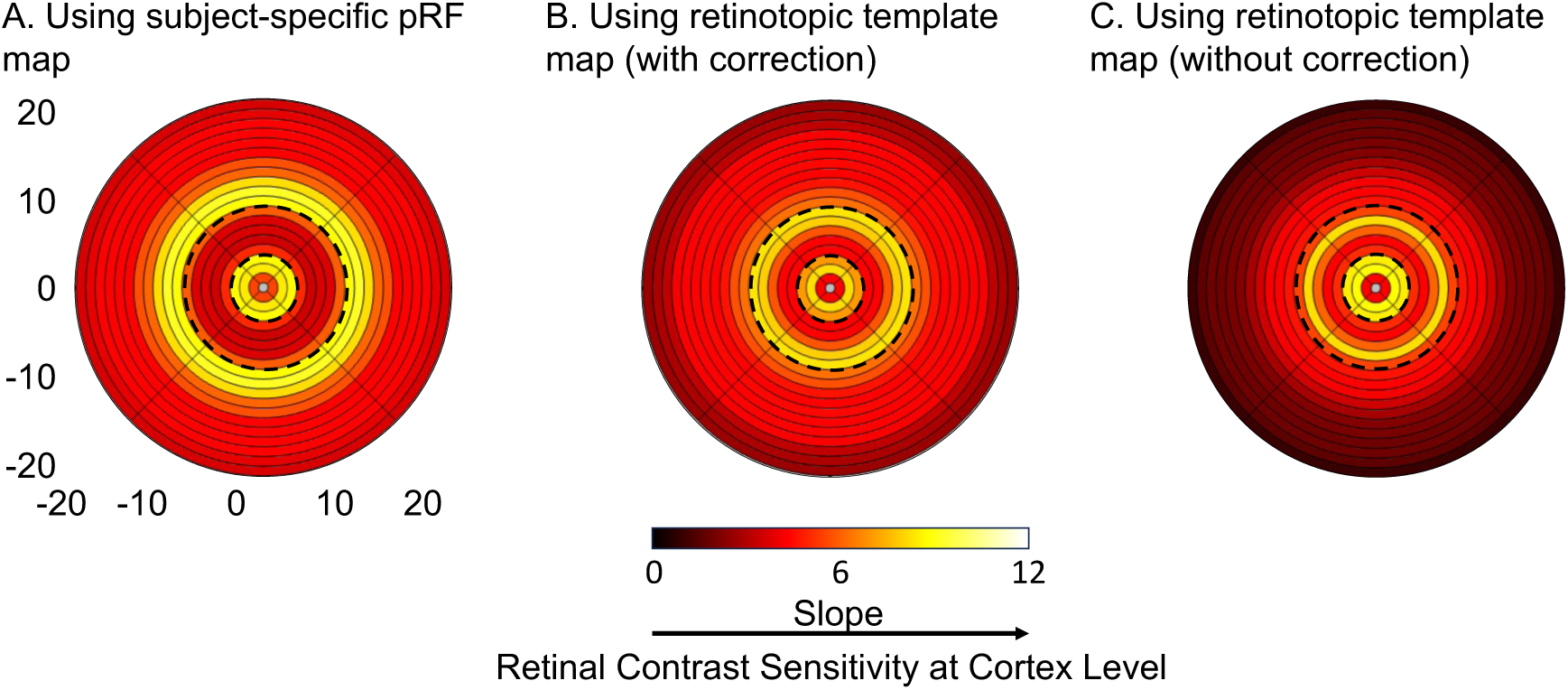
Effect of correcting eccentricity distribution in retinotopic templates on cortical contrast sensitivity maps. Heatmaps show slope values back projected into visual space, with each concentric circle representing a 1° eccentricity bin. A: Back projection was performed using subject-specific pRF map, B: Using Benson atlas with eccentricity distribution corrected using the H&H model. C: Using Benson atlas without correction (original template). Color scale indicates slope values reflecting cortical contrast sensitivity (brighter = larger slopes).

